# Cepo uncovers cell identity through differential stability

**DOI:** 10.1101/2021.01.10.426138

**Authors:** Hani Jieun Kim, Kevin Wang, Carissa Chen, Yingxin Lin, Patrick PL Tam, David M Lin, Jean YH Yang, Pengyi Yang

## Abstract

We present Cepo, a method to generate cell-type-specific gene statistics of differentially stable genes from single-cell RNA-sequencing (scRNA-seq) data to define cell identity. Cepo outperforms current methods in assigning cell identity and enhances several cell identification applications such as cell-type characterisation, spatial mapping of single cells, and lineage inference of single cells.

---

Defining cell identity is fundamental to understand the cellular heterogeneity in the population and the cell-type-specific response to environmental signals and experimental perturbations. Exploring cell identity has been enabled by rapid technological advances in genome-wide profiling of the molecular content in single cells^1–3^. This comprehensive lens into the molecular properties unique to each cell type allows defining and predicting cell identities in ways that were previously not feasible using data generated by bulk/population analytics technologies. The most widely used method to define genes associated with cell identity is differential expression (DE)^4,5^. Despite the advances in high-throughput scRNA-seq data that would provide unprecedented resolution of cell identity, none of the current approaches^6^ has been evaluated systematically for their attribute and fidelity for defining cell identity genes (CIGs) from scRNA-seq dataset of millions of cells and consisting of hundreds of cell types.

We developed Cepo (refers to “cell” in Korean), a method to retrieve genes defining cell identity from scRNA-seq data. We propose a biologically motivated metric, differential stability (DS), to identify cell-type specific genes on the premise that stable gene expression is a key indicator of cell identity. Our hypothesis implies that genes marking a cell type should be (1) expressed and (2) stable in its expression level relative to other cell types. We translate the criteria into a computational framework where, using pre-defined celltype labels, we compute cell-type-specific statistics to prioritise genes that are DS against other cell types in all cell-type pair comparisons (Fig. 1a, Online Methods: Cepo implementation).

**Figure 1.**
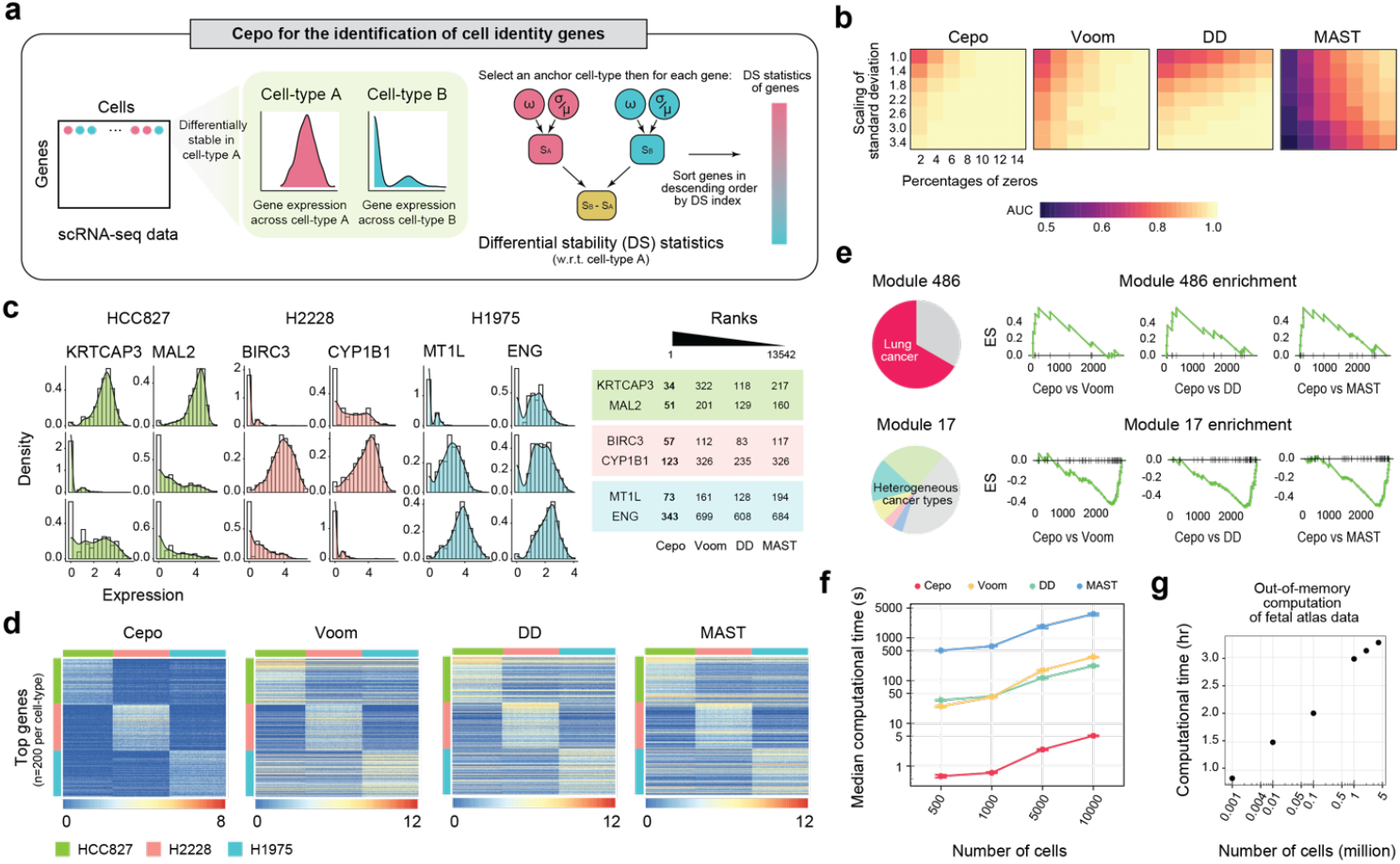
Cepo uncovers differentially stable genes in synthetic and experimental single-cell RNA-sequencing datasets. **(a)** A schematic of Cepo to define the differential stability (DS) statistics. **(b)** AUC of Cepo and other differential analysis methods, Voom^7^, differential distribution (DD), and MAST^9^, in simulated scRNA-seq datasets containing 500 differentially stable genes between two cell types. The degree of DS was modulated by increasing the percentage of zeros (column) and/or by increasing the scaling of the standard deviation of gene expression (row). **(c)** Distribution of expression of example DS genes identified for each cell type from a scRNA-seq dataset of three lung adenocarcinoma cell lines^10^. The ranking of each gene in their respective cell types of interest by each method are tabulated. A lower rank denotes higher prioritization of the gene. **(d)** Expression of top 200 genes identified for each of the three lung adenocarcinoma cell lines by the differential analysis methods. Columns correspond to cells and rows correspond to individual genes. For each cell type, the genes are ordered by decreasing differential statistics. **(e)** Gene set enrichment plot of Cancer Modules^11^ 486 and 17 on the relative ranks of the top 3000 genes from Cepo and each of the differential analysis method (y-axes denote enrichment scores and x-axes denote the rank difference). **(f)** The median computation time to run each method in seconds for n = 20 independent simulations across varying number of cells and 20,000 genes are shown. Error bars denote the lower and upper quantiles. **(g)** Computation time of Cepo on the fetal atlas data consisting of approximately 4 million single cells^12^.

We performed a comprehensive benchmarking of Cepo with several differential analysis methods^6–9^ using both simulated and experimental scRNA-seq datasets in a range of biological systems that require knowledge of cell identity. We tested the accuracy and efficacy of the method to detect subtle changes in stability against simulated DS genes with varying stability (Supplementary Fig. 1 and Online Methods). We show that Cepo, followed by Voom, can identify simulated DS genes with the highest accuracy relative to other methods (Fig. 1b and Supplementary Fig. 2, 3).

To determine the effectiveness of Cepo to detect DS genes, we applied Cepo on the CellBench data consisting of single-cell expression profiles from three human lung adenocarcinoma cell lines^10^. Evaluation of the global ranks of the differential statistics generated from each method and the two components of Cepo, relative CV (rCV) and relative proportion of zeros (rzprop; see Online Methods), demonstrated a strong correlation between the differential analysis methods and also with rCV (Supplementary Fig. 4, 5). In agreement with the simulation results, among the benchmarked methods, Voom demonstrated the strongest correlation with Cepo and rCV. Genes identified by Cepo showed stronger differential stability between cell types when compared to other competing methods (Fig. 1c, d and Supplementary Fig. 6). Consistent with prioritisation of differences in mean gene expression by other differential analysis methods, genes specifically prioritised in these methods showed less DS between cell types (Supplementary Fig. 6–7). To determine whether the DS genes prioritised by Cepo enhance the assignment of cell identities, we performed gene set enrichment against Cancer Module database^11^ on the relative scores of the top ranked genes between Cepo and other differential analysis methods. When comparing the genes identified from lung adenocarcinoma (H2228) cells using Cepo and other methods, we observed a strong enrichment of lung cancer and the depletion of other heterogeneous cancer types from the Cancer Module Database (Fig. 1e and Supplementary Fig. 8). These results demonstrate the ability of Cepo to prioritise relevant cell identity gene sets and de-prioritise the irrelevant ones.

As the number of cells covered by the dataset increases, computational speed becomes a key factor in method selection. Cepo is computationally fast, requiring only seconds to analyse datasets with tens of thousands of single cells (Fig. 1f). Whilst dedicated single-cell methods (such as MAST) have been shown to be much slower than bulk methods (such as Voom)^5^, we demonstrate that Cepo is fastest in computation (Fig. 1f) and scales up effectively when applied to dataset containing millions of single cells^12^ (Fig. 1g).

High throughput single-cell data is requisite for the characterisation of rare cell types. It is therefore imperative to develop methods that can detect genes associated with cell identity in cell types that make up a minor proportion of the population. To address this challenge, we devised an experiment evaluating the reproducibility of CIG detection in rare cell types using data subsampling (Online Methods). We found that whilst the global statistics was fairly reproducible across all methods (Fig. 2a and Supplementary Fig. 9), identification of the top CIGs was most consistent by Cepo (Fig. 2b). Besides the subsampling analysis, we also observed strong concordance of Cepo-derived statistics across technologies and batches in scRNA-seq data (Supplementary Fig. 10).

**Figure 2.**
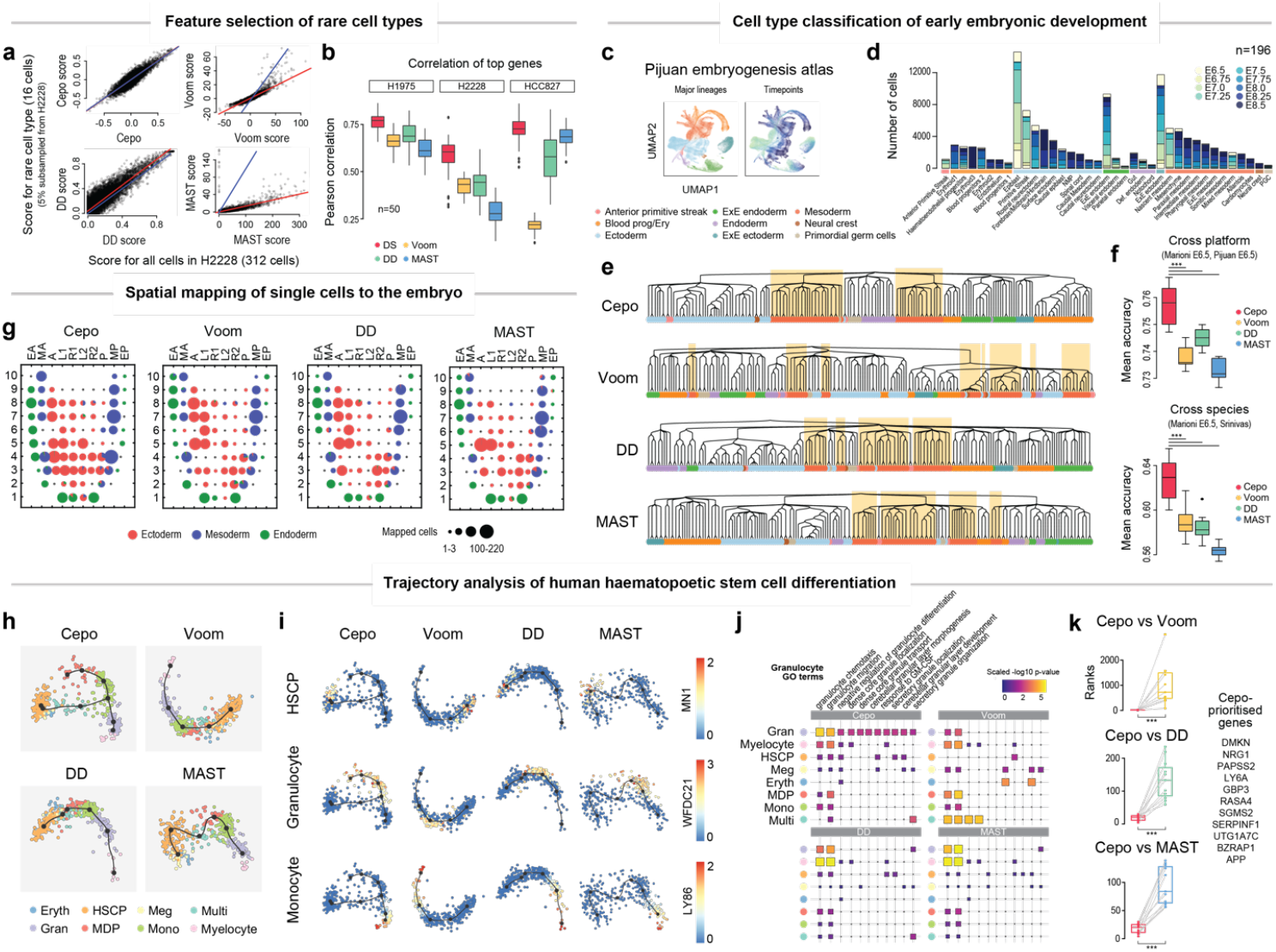
Cepo effectively retrieves cell identity genes and enhances interpretation of diverse single-cell applications associated with cell identity. **(a)** Scatter plot of differential analysis scores from the full dataset (x-axis) and the rare cell type dataset (y-axis) generated by subsampling 5% of the total number of cells from cell type H2228. The red line denotes the best line of fit, and the blue line denotes x=y. **(b)** Boxplot of Pearson’s correlation between ground-truth and rare cell-type scores of the top 40 genes repeated 50 times. **(c)** UMAP visualization of the embryogenesis atlas scRNA-seq data highlighted by 9 major lineages or cell types (left) and embryonic time-points (right). **(d)** Barplot of cell numbers for each cell type. Each bar is sub-sectioned by the proportion of cells found in each embryonic time point, giving a total of 196 samples. The bars are grouped into the 9 major lineages or cell types and ordered by decreasing cell count. **(e)** Clustering of 196 samples by HOPACH trees. The terminal nodes are coloured by the major lineage/cell-type labels. Sections of the tree have been shaded to highlight clusters of the mesoderm lineage samples. **(f)** Cross-platform classification of E6.5 single cells from the mouse 10X dataset^14^ using a kNN trained on the mouse Smart-seq2 dataset^13^ (upper panel), and crossspecies classification of Carnegie Stage 7 single cells from the human Smart-seq2 dataset^15^ using the kNN trained on E6.5 single cells from the mouse Smart-seq2 dataset^14^ (lower panel). **P < 0.01; ***P < 0.001, two-sided paired Wilcoxon test. **(g)** Spatial mapping of E7.5 single cells from the Smart-seq2 dataset^16^ onto the E7.5 embryo (n = 83 spatial locations). Each dot denotes a pie chart that shows the proportion of cell types mapped to the location. **(h)** Trajectory inference of hematopoietic stem cell differentiation dataset using multi-spanning tree and genes identified by the differential analysis methods. **(i)** Trajectories are highlighted by expression of key marker genes. **(j)** Gene set over-representation analysis of GO terms associated with granulocytes. **(k)** Paired boxplots of ranked differential statistics of select monocyte-associated genes between Cepo and other differential analysis methods. **P < 0.01; ***P < 0.001, two-sided paired Wilcoxon test.

We reasoned that the integration of Cepo with diverse applications of single-cell analysis for the investigation of the spectrum of cell types would enhance retrieval of cell identity. We first analysed a high-throughput embryogenesis atlas data^13^ that cover multiple lineages, complex cell types, and time points (Fig. 2c-d) to test the ability of differential analysis methods to cluster like cell types at different time points. We expect that global statistics between cell types of the same lineage should be more similar than those from more distant lineages. Indeed, visualisation of the pairwise correlation of differential statistics revealed stronger correlation with cell types from the same lineages, implying that the Cepo statistics may be more informative of cell identities than other methods (Supplementary Fig. 11–14). Furthermore, clustering analysis revealed that Cepo was able to group the major lineage/cell types more effectively than the other differential methods, as exemplified by the clustering of mesoderm lineage samples (Fig. 2e and Supplementary Fig. 15). Expression of genes specifically prioritised by Cepo and displayed in UMAP confirmed the differential stability of these genes was closely associated with cell identity (Supplementary Fig. 16). Moreover, we showed that Cepo-defined CIGs lead to more accurate classification of cell-types using scRNA-seq data generated by different technologies^14^ and for different species^15^ (Fig. 2f).

Factors influencing cell identity include the spatial location and neighbourhood of the cell. We considered whether CIGs identified by Cepo include genes associated with the spatial environment on the premise that factors influencing cell identity may include the environment in which the cell resides^3^. When applied Cepo to the mapping of single cells from E6.5 and E7.5 mouse embryos^14^ to tissue domains in embryos of the corresponding timepoints^16^, single cells were regionalised to tighter locations in the germ layers (Fig. 2g and Supplementary Fig. 17a) with greater homogeneity of cell types in each location than the other differential analysis methods (Supplementary Fig. 17b). We further confirmed the gene expression of spatial markers were in agreement with prior biological knowledge (Supplementary Fig. 17c), highlighting the attribute of Cepo to assign high-confidence cell identities at defined locations.

Demonstration of the utility of Cepo’s in identifying cell types led us to query the relevance of Cepo-identified DS genes to recapitulate the underlying biology in stem cell differentiation. Current trajectory inference tools commonly rely on the judiciously selected genes, as inclusion of all genes may mask the biology and introduce noise in data analysis. To test this, we tested whether the Cepo-selected CIGs were able to recapitulate the branching of the granulocyte and monocyte lineages^17,18^ using a scRNA-seq dataset profiling hematopoietic stem cell differentiation^19^ (Supplementary Fig. 18a). We found that trajectory inference on genes selected using other differential analysis methods was unable to recapitulate the branching of the two lineages (Fig. 2h-i). In contrast, genes selected by Cepo uncovered the underlying structure of the data, resulting in revealing the branching of the monocyte and granulocyte lineages (Fig. 2h-i). We corroborated this finding by running the trajectory analysis with genes that are exclusive to each method as well as genes common to all four methods by showing that use of Cepo-exclusive genes alone is sufficient to retrieve the bifurcation event (Supplementary Fig. 18b-c). As a demonstration of robustness, Cepo consistently captured the bifurcation of the lineage tree across multiple trajectory inference methods (Supplementary Fig. 19). Gene set over-representation analysis for granulocytes revealed enrichment of many terms associated with the granulocyte function (Fig. 2j). Similarly, in other cell types, we found that Cepo prioritised genes associated with cell identity more effectively than other methods (Fig. 2k and Supplementary Fig. 20; see Online Methods for references). For instance, in monocytes, Cepo identified genes such as neuregulin-1 (NRG-1), an endogenous activator of the NRG-1/ErbB pathway, which regulates macrophage and monocyte function^20^ (Fig. 2k).

Cepo is a biologically driven method to uncover cell identity and cell-fate decisions (Fig. 1e and Fig. 2e-k). Inherent to the design of Cepo is the ability to scale to datasets containing millions of cells on a laptop (Fig 1g). Cepo is compatible with 3’ and full-length sequencing protocols (Supplementary Fig. 10). In contrast to DE methods, Cepo incorporates information from both highly and lowly expressed genes and prioritises genes more efficiently than other methods that rely on more highly or differentially expressed genes (Supplementary Fig. 6). As a method for CIG identification, Cepo will facilitate the mining of the growing resource of single-cell data and realise the potential of single-cell analytics technologies to pinpoint cell identities that are relevant to the cellular phenotype.

## Online Methods

### Datasets

The following datasets were used to evaluate Cepo and demonstrate its use in extracting cell identity from diverse sources of single-cell data and a spatial embryo dataset generated using geographical position sequencing (GEO-seq). All the datasets used in this study are publicly available.

- **CellBench data** The “CellBench data” collection^1^ was downloaded from https://github.com/LuyiTian/sc_mixology/. We use two scRNA-seq datasets from five human lung adenocarcinoma cell lines HCC827, H1975, H2228, H838, and A549, which were cultured separately. The first dataset contains the first three cell lines (i.e., HCC827, H1975, H2228) and the second the entire set. To generate the libraries, single cells from each cell line were mixed in equal proportions, and then the data were generated using 2-3 different protocols: CEL-seq2, Drop-seq and 10x Genomics Chromium. The resulting read counts were normalized using the scran package^2^.
- **Embryogenesis atlas data** The “Embryogenesis atlas data”^3^, which profiles 48 hours of mouse embryonic development, was downloaded from https://github.com/MarioniLab/EmbryoTimecourse2018. Staged mouse embryos at 9 different time points were collected, prepared into single-suspensions, and then scRNA-seq libraries were generated using the 10x Genomics Chromium system. The resulting read counts were log-transformed and size-factor adjusted.
- **Gastrulation data** The parsed “Gastrulation data”^4^, sequenced using scNMT-seq^5^, was downloaded from the link provided in https://github.com/rargelaguet/scnmt_gastrulation. The resulting read counts were log-transformed and size-factor adjusted.
- **Human embryo data** The processed “Gastrulation data”^6^ was downloaded from http://www.human-gastrula.net. scRNA-seq of human CS7 embryo was generated using Smart-seq2 protocol^7^. The data were normalized using ‘quickcluster’ and ‘normalize’ functions from the scran package^2^. This was followed by pseudocount addition of 1 and natural-log transformation of the count matrix.
- **Haematopoietic stem cell differentiation data** The “HSC differentiation data”^8^ was downloaded from https://cytotrace.stanford.edu/. Bone marrow cells were harvested from euthanized mice, enriched, and FACS-sorted for generation of singlecell libraries using the C1 Single-Cell Auto Prep System (Fluidigm). The resulting count matrix was TPM/FPKM normalised.
- **Fetal tissue atlas data** The “Fetal tissue atlas data”^9^ was downloaded from NCBI Gene Expression Omnibus under accession number GSE156793. Approximately 4 million single-cell transcriptomes were generated using the sci-RNA-seq3 protocol^10^ from 121 human fetal samples, representing 15 organs.
- **GEO-seq spatial embryo data** The “spatial embryo data”^11^ was downloaded from NCBI Gene Expression Omnibus under accession number GSE120963. The spatial data was generated using the protocol described in Peng et al.^12^ from mouse embryos dissected at 5 embryonic time points.

For all datasets, only cells that passed the quality control of the original publication and assigned cell types were included. If only the count matrix was supplied, we performed size factor normalisation of the raw count matrices and then generated the log-transformed gene expression matrices using the ‘logNormCounts’ function from the scater R package^13^. We otherwise used the fully processed matrix provided in the above links. Original cell-type labels were used as indicated in the original publication and doublets were removed according to the labels where provided.

### Cepo implementation

A fundamental goal of single-cell biology is to identify how experimental manipulation or environmental signals affect individual cell-types within a population. Current approaches rely on differential gene expression (DE), which has been the go-to method to find genes associated with cell identity^14,15^. However, these methods often rely on the differences in mean gene expression. Whilst differences in gene expression level may be a key discriminating factor between cell types, how well these differences reflect the identity of the cell remains unclear. In this study, we propose “differential stability (DS)” as a new metric to define cell identity genes (CIGs) under the premise that stable gene expression is a key indicator of cell identity. DS prioritises genes that are (i) expressed and (ii) stable in its expression level relative to other cell types. By contrast, DE methods would prioritise genes that may be stably expressed in the cell type of interest and also others, as long as they are differentially expressed. We argue that for a gene to accurately represent a cell’s identity, stable expression of the gene in that cell type is required (or the lack of in other cell types).

We recently proposed a computational framework for characterising stability of gene expression across single cells, demonstrating the concept of housekeeping genes to identify genes that are characteristic of individual cell types within a population^17^. Extending on this, here we assess DS of genes *i*(*i* = 1,…, *G*) in each cell type *j*(*j* = 1,…, *M*) by using a combination of stability features including proportion of zeros ***ω**_j_*, mean expression level ***μ**_j_*, and variation of expression 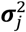. Specifically, we first calculate a vector of stability score of genes in cell type *j*(*j* = 1,…, *M*) as the average of normalised ranks of ***ω**_j_* and coefficient of variation (CV):

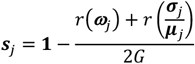

where *r*(.) is the function that generates the ranks based on a vector of input values. This allows us to quantify the stability of each gene *i* in cell type *j* where the smaller the proportion of zeros and CV of the gene, the larger the value of s_ij_. The DS score of genes for the *j*th cell type is then defined as:

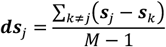

Sorting the values of ***ds**_j_* in decreasing order allows the most differentially stable genes to be ranked at the top of the list for the yth cell type.

### Differential analysis methods

We performed other differential analyses of gene expression using the Model-based Analysis of Single-cell Transcriptomics (MAST) method^18^, Voom and Limma-trend (Limma) implemented in limma R package^19^, edgeR^20^, differential distribution (DD) based on the Kolmogorov-Smirnov test (KS-test)^21^, as well as simple *t*-test and Wilcoxon test. For this study, we chose DE methods that were recently demonstrated to have high performance in a benchmarking study on scRNA-seq data^15^. We used the Benjamini-Hochberg^22^ method for the adjustment of *p*-values. All functions used default parameters. For the CellBench benchmarking study, we ran edgeR quasi-likelihood (QLF) test and included the cellular detection rate (i.e. the fraction of detected genes per cell) as a covariate on TMM-normalised count matrix, as recommended in Soneson et al.^15^. The size-factor normalized and log-transformed expression matrix was used for other differential analysis methods unless otherwise stated.

To compare the differential analysis statistics from the above methods against differential stability, we compute the two components of Cepo—coefficient of variation and proportion of zeros—individually for evaluation purposes only. We compute relative CV (rCV) for the *i*th gene in the *j*th cell type is below:

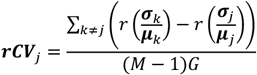

The relative proportion of zeros (rzprop) value can be defined identical to rCV by replacing the CV calculation with a calculation for the proportion of zeroes.

### Simulations

We perform a formal assessment on the ability of the benchmarked differential analysis methods to detect DS genes using simulation. We first take the CellBench data^1^ and subset it to only contain the H1975 and H2228 cell types. We then use the Splatter package^23^ to both (1) estimate distributional parameters in this dataset and (2) simulate a new single-cell (log-transformed counts) data using the estimated parameters by fixing the number of genes to 10,000.

We simulate DS genes with distributional differences between two randomly assigned cell types, named A and B. We first calculate *p_i_*, defined as the product between the mean and proportion of non-zero expression values for a given gene, *i*. We then define 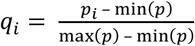, which scales all *p_i_*’s to be in the interval [0, 1]. We then sample 100 genes proportional to the weights *w_i_*, where *w_i_* is defined as *q_i_* if *q_i_* > 0.2 and 0 otherwise. This allows the sampling of genes with lower proportion of zeros as candidates for DS genes before we induce distributional differences. We then introduce DS genes in the two cell types (where cell type A is regarded as the cell type of interest) by performing the following steps for each DS gene:

1. For all non-zero expression values, we calculate the mean, *m*, and standard deviation, *s*.
2. Generate *X* = (*x*_1_,…, *x_nz_*) where *nz* is the number of cells with non-zero expression value for that gene. Each *x_c_* is an independently generated normal random variable with mean *m* and standard deviation *αs*, where *α* is a scaling factor equally spaced between 1 and 4. Any non-positive value of *x_c_* is then set to zero to induce sparsity.
3. We take the remaining non-zero values from step 2 and replace *β* percent of these values with zero. This *β* is an additional proportion of zeroes and we choose it to be equally spaced between 0% and 14%.

This procedure is an attempt to: (1) increase the spread of the simulated DS gene through the parameter *α*, and; (2) increase the sparsity in cell type B through the thresholding in step 2 and the *β* parameter in step 3. In Supplementary Fig. 1, the values of *α* and *β* as rows and columns, respectively.

Given that we have 100 DS genes out of a total of 10,000 genes, we then calculate the area under the receiver operating characteristic curve (AUC-ROC) based on the ranks of the genes by all four differential statistics using the yardstick package^24^. The heatmaps in Fig. 1b and Supplementary Fig. 3 show the AUC-ROC value averaged across 100 such simulation for each of all differential methods.

### Computational time benchmarking

Fixing the number of cell types to 3 and the number of genes to 20,000, we simulate matrices with 500, 1000, 5000 and 10,000 cells. These matrices are then evaluated by Cepo, Voom, DD and MAST using R with 20 repetitions. The computational times are presented in Fig. 1f.

To highlight the performance of Cepo in the computation of data with very large number of cells, we used Cepo to perform out-of-memory computation of a human fetal tissue atlas comprising over 4 million singlecell transcriptomes^9^. In particular, Cepo handles matrices in the class of ‘DelayedArray’ using ‘HDF5Array’^25^, which allows out-of-memory computation. Although this would slow down computational speed compared to in-memory computation, out-of-memory computation has a desirable advantage of enabling computations on data that is much larger than what the computer RAM can fit at once. This eliminates the issue of memory wall when performing Cepo on ultra-high-throughput data. All evaluation was performed on a research server with dual Intel(R) Xeon(R) Gold 6148 Processor (40 total cores, 768 GB total memory) and dual RTX2080TI GPUs.

### Rare cell-type and reproducibility analysis

Large-scale single-cell technologies have enabled discovery of rare cell subpopulations^26^. To assess the capacity of Cepo to generate consistent results for rare cell types, we devised an experiment to evaluate the reproducibility of CIG detection in subsampled data. For each cell type, we performed subsampling of the cell type to 5% of its original size, whilst leaving the sample size for the other cell types as they are. We performed this subsampling 50 times for each differential analysis method and compared the results to those generated from the full dataset. To evaluate the concordance of the highly ranked genes, we used Pearson’s correlation to calculate the concordance of the top 40 genes, identified using full sample data, for the 50 subsampling runs and for each cell type.

To test the reproducibility of the Cepo with regards to both single-cell sequencing technology and batch, we compared the differential stability statistics from scRNA-seq data generated from different sequencing technologies and batch. We used the CellBench datasets^27^, which are particularly amenable for such benchmarking analyses. The CellBench scRNA-seq mixology experimental design enables both cross-technology (CEL-seq2, 10x Genomics Chromium, and Drop-seq) and cross-batch analysis of three cell types, H2228, H1975, and HCC827. The second batch was sequenced on the 10x Genomics Chromium platform and contained two more cell types (H838 and A549).

### Clustering of differential analysis statistics

We performed hierarchical ordered partitioning and collapsing hybrid (HOPACH) clustering^28^ to evaluate sub-cluster relatedness. We used the distance matrix generated from the rank-transformed differential analysis statistics. For the HOPACH clustering, we set a value of 10 for ‘kmax’, which controls the maximum number of children at each node, whilst keeping the default settings for all other parameters. Visualisation of the correlation between the differential analysis statistics was generated using the pheatmap R package^29^ on the correlation matrix. Spearman’s correlation was used to calculate the correlation between the statistics.

### Classification of single cells

We assessed the utility of genes identified using Cepo, MAST, Voom, and DD on classifying single cells to their respective cell types across different scRNA-seq profiling technologies and species. Specifically, we first trained a k-nearest neighbour (kNN) classifier to classify cells to the three cell types of Visceral Endoderm, Epiblast, and Primitive Streak from the mouse embryos at Embryonic day 6.5 (E6.5) profiled using Smart-seq2^4^ and 10x Genomics Chromium^3^ scRNA-seq technologies. Next, we trained the kNN using the cells from mouse embryos at E6.5^4^ and classified the cells profiled from human embryos from the corresponding developmental stage (Carnegie stage 7)^6^. The Pearson’s correlation-based similarity metric and the k value of 10 were used for kNN classifiers given their good performance across a large number of datasets in our previous work^30^ and the union of top 25 genes selected from each of all cell types by each of the four methods were used as learning features of kNN. To assess the classification accuracy, we performed cell type-stratified subsampling using 80% of the cells from each cell type and calculated the average accuracy across the three cell types. We repeated this subsampling procedure 10 times to estimate the performance variability.

### Spatial mapping of single cells to spatial embryo data

We used the single cells profiled using Smart-seq2 scRNA-seq from the mouse embryos at E6.5 and E7.5^4^ and the corresponding spatial transcriptomes of mouse embryos at these two development stages^11^ to assess the utility of genes selected from Cepo, Voom, DD, and MAST on mapping single cells to their spatial locations. In particular, we first obtained the union of the top 100 genes selected from each of the three cell types (Visceral Endoderm, Epiblast, and Primitive Streak for E6.5; Ectoderm, Endoderm, and Mesoderm for E7.5) and then assigned each single cell to the best matched spatial location based on Spearman’s correlation of the expression of the selected genes. To quantify the performance of the four differential analysis methods (i.e., Cepo, Voom, DD, and MAST) on mapping single cells to spatial transcriptomes of the embryos, we calculated the purity of spatial locations based on the assigned single cells. Specifically, we define the purity of a spatial location *l*(*l* = 1,…, *L*) as *purity_l_* = max (*p*_1_, *p*_2_,…, *p_M_*), where 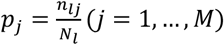 is the percentage of cell type *j* at that spatial location of the embryo, *n_lj_* is the number of cells in that cell type assigned to *l* and *N_l_* is the total number of cells assigned to *l*. The overall purity of single-cell mapping to spatial transcriptomes can then be quantified as 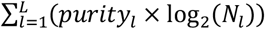, where the term log_2_(*N_l_*) accounts for the number of cells assigned to each spatial location.

### Trajectory analysis

To evaluate the relevance of Cepo-identified DS genes to recapitulate the underlying biology in stem cell differentiation, we performed trajectory analysis using the marker genes defined by the differential analysis methods. To perform the trajectory analyses, we employed the dynverse framework^31^, which facilitate benchmarking of trajectory inference tools. We inferred trajectories using the following tools: 1) the multispanning tree (MST) approach. MST requires the dimensionality reduced matrix as input, for which we used multi-dimensional scaling; 2) Slingshot^32^; 3) projected Monocle2^33^; 4) Monocle2^33^ with DDR-TREE, a scalable reversed graph embedding algorithm for dimensionality reduction; 5) CellTree^34^ using the maptpx method^35^ for latent Dirichlet allocation (LDA); 6) Slice; 7) CellTree^34^ using the variational expectation-maximisation (vem) method for LDA; and 8) Monocle2^33^ with ICA for dimensionality reduction. In each case, the default settings were used. The top 200 genes with the highest differential statistics from each cell type were pooled and used to build the trajectories. The original cell type labels were used where cluster assignments were required as input. To determine whether the resulting trajectories has recapitulated the correct bifurcation event, we first calculated the pseudotime of differentiation on the basis of the inferred trajectories, setting a random cell in the HSCP cell type as “root” cell. We then considered a tree or bifurcation as a trajectory with a correct bifurcation event when the terminal cell types included Myelocytes or MDPs. The terminal cell types were defined as cells found between 0.7 and 1 of the pseudotime, where 1 denotes the terminal cell and 0 the root cell.

To investigate whether the genes specifically identified by Cepo are responsible for the observed bifurcation, we repeated the above trajectory analysis on a refined gene set wherein genes that are specifically identified by each method were used. Briefly, the top 200 genes with the highest differential statistics were selected from each cell type as before. Before pooling the genes together, we excluded any genes that were not part of (1) the common gene set of genes identified by all four differential analysis methods (Cepo, Voom, DD, and MAST) and (2) the method-specific genes that were specifically identified by the method-of-interest. These method-specific gene sets were then used in the trajectory analysis under the same conditions as above.

### Functional enrichment analyses

Gene set enrichment analysis was performed using the fgsea R package^36^ using pathway annotations from the MSigDB database^37^ and in particular the Cancer Gene Co-expression Modules^38^. We use the relative ranks between the differential analysis statistics to assess the relative enrichment of gene sets between two methods. The relative enrichment analysis was performed on the top 3000 genes. Amongst the most significantly enriched cancer modules were 486 and 17. Each cancer module in the Cancer Gene Coexpression Modules are characterised by up- and down-regulated clinical annotations. Because each differential analysis method prioritises genes that demonstrate increased differential stability, expression or distribution in the cell type of interest with respect to other cell types, we considered only the up-regulated clinical annotations when characterising the cancer modules. The cancer types associated with the up-regulated clinical annotations in Cancer module 486 include lung (6) and liver (3) cancers. Those associated with the up-regulated clinical annotations in Cancer Module 17 are liver cancer (11), stimulated PBMCs (6), stimulated immune (4), breast cancer (2), B lymphoma (1), and neuro tumours (1). The number inside the brackets denotes the total number of annotations for each cancer type. Lastly, the enrichment analyses were visualised using the ‘plotEnrichment’ function.

Pathway over-representation analyses were performed using the ‘enrichGO’ function in ClusterProfiler R package^39^. The biological process ontologies were used, and the P values were adjusted for multiple testing using Benjamini-Hochberg FDR correction at *α* = 0.05. To perform the over-representation analysis, we used 100 genes with the highest statistics.

## Data availability

The count or expression matrices used in this study can be found at the links provided at the “Datasets” section of the article or the raw data can be found at Gene Expression Omnibus under the accession numbers described in “Datasets”.

## Code availability

The source code used to perform the analyses presented here is available at https://github.com/PYangLab/CepoMauscript. Cepo R package and the detailed vignette are available at https://github.com/PYangLab/Cepo.

## Author contributions

P.Y. and H.J.K. conceived the study with input J.Y.H.Y. and D.M.L.; H.J.K. and K.W. developed the method and software with input from P.Y.; H. J.K., P.Y., and K.W. performed data analyses with input from C.C. and Y.L.; H.J.K., P.Y., K.W. and J.Y.H.Y. interpreted the results with input from P.P.L.T.; H.J.K., P.Y. and K.W. wrote the manuscript with input from J.Y.H.Y.; All authors read and approved the final version of the manuscript.

## Conflict of interest

The authors declare no competing interests.

## Acknowledgments

The authors thank all their colleagues, particularly at the School of Mathematics and Statistics, The University of Sydney, and Sydney Precision Bioinformatics Alliance for their support and intellectual engagement. This work is supported by an Australian Research Council (ARC)/Discovery Early Career Researcher Award (DE170100759) and a National Health and Medical Research Council (NHMRC) Investigator Grant (1173469) to P.Y., an Australian Research Council Discovery Project grant (DP170100654) to P.Y. and J.Y.H.Y., and an Australian Research Council (ARC) Postgraduate Research Scholarship and Children’s Medical Research Institute Postgraduate Scholarship to H.J.K.

---

**Supplementary Figure 1.**
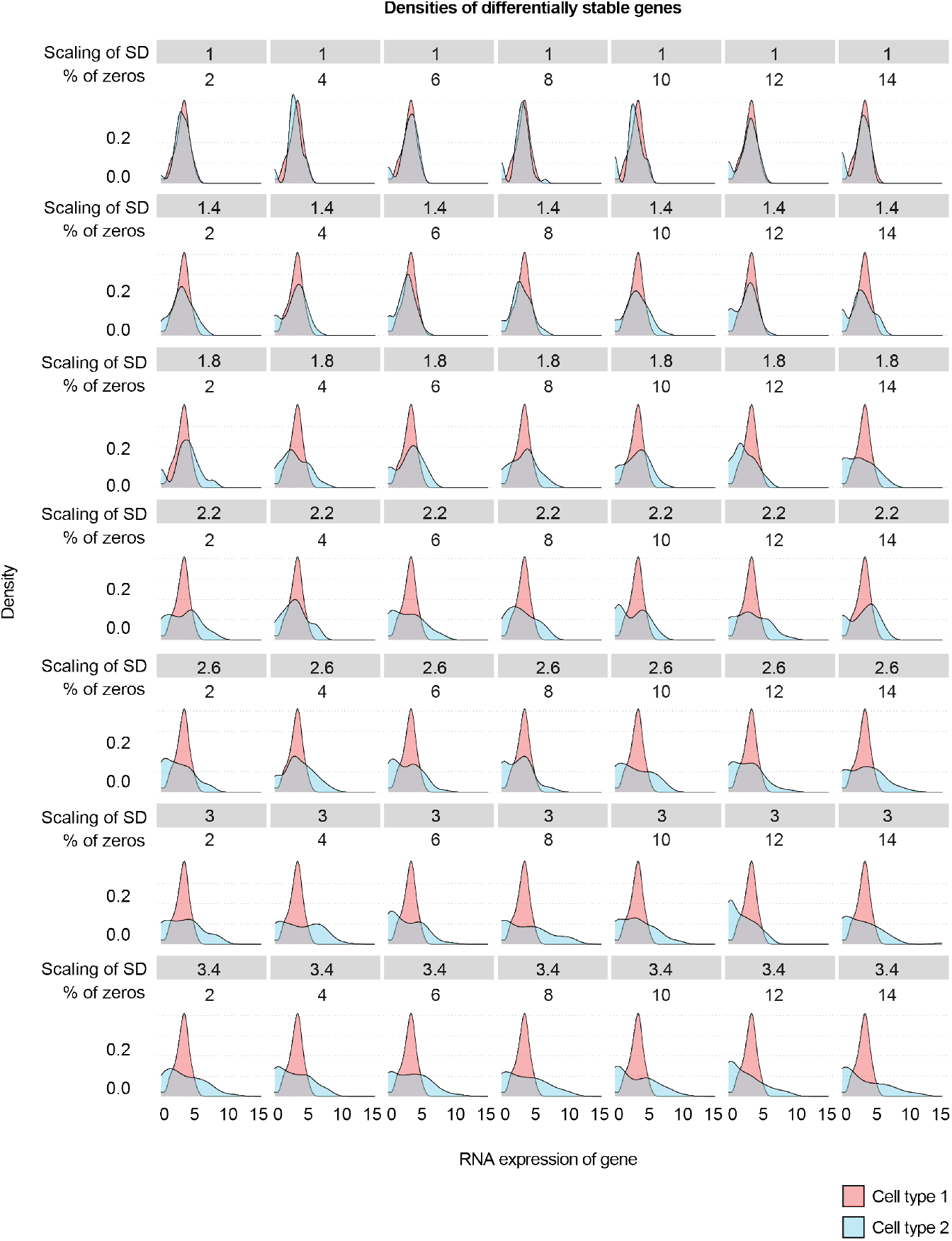
Simulation of differential stability genes. In each simulated scRNA-seq dataset, DS genes in Cell type 1 were simulated by varying the proportion of zeros (columns) and standard deviation (rows) of gene expression in Cell type 2. These two parameters correspond to the *β* and *β* parameter in Online Methods. The densities of gene expression for one exemplar gene are shown above.

**Supplementary Figure 2.**
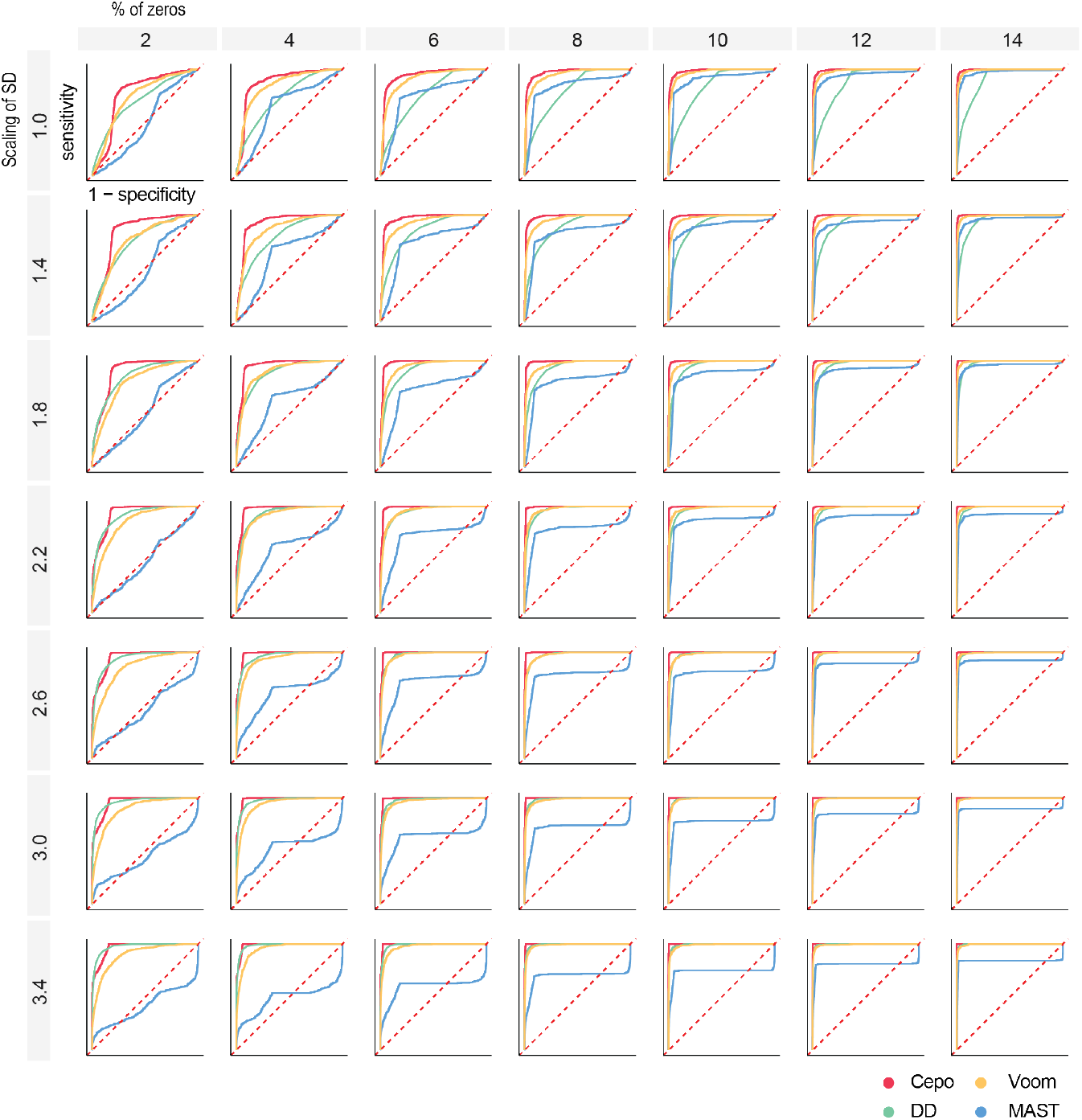
ROC curves of differential stability gene detection. The ROC curves were used to assess the capacity of each method to detect DS genes between two cell types. The curves are colour-coded by method. The 1-specificity and sensitivity of DS gene detection are plotted on the x- and y-axes, respectively. The dashed diagonal lines denote the identity (y = x) lines.

**Supplementary Figure 3.**
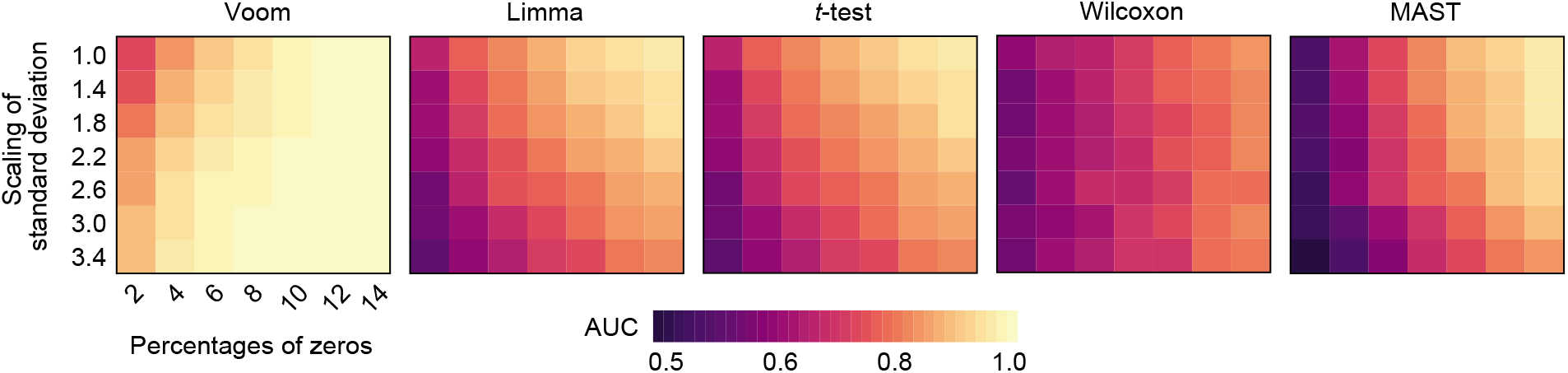
Heatmap of AUCs from each differential analysis method. The area under the ROC curve (AUC) denotes the accuracy of DS gene detection for simulated scRNA-seq datasets are visualised as a heatmap for each differential analysis method.

**Supplementary Figure 4.**
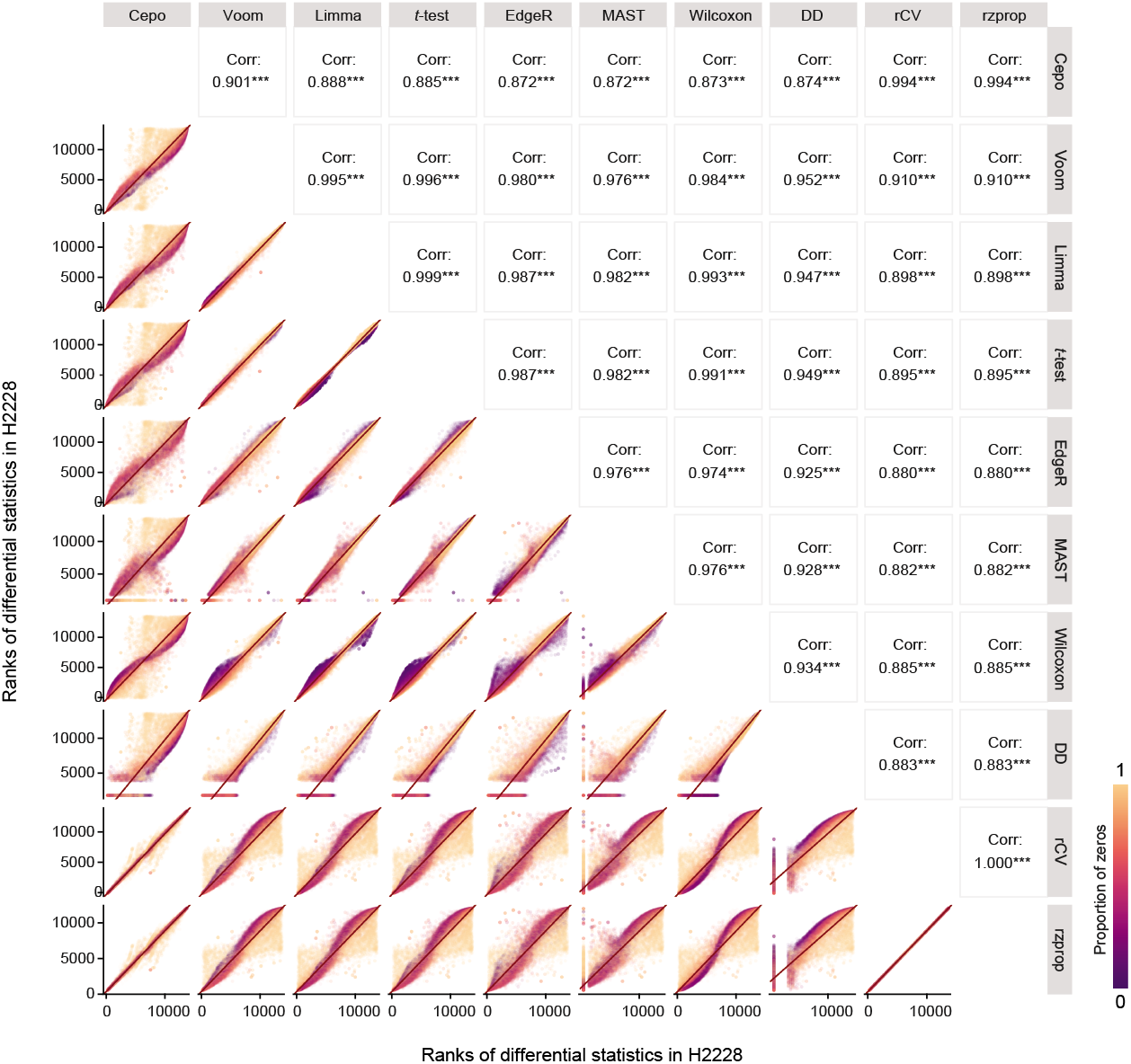
Pairwise scatter plot of ranked differential statistics for H2228. The statistics generated for H2228 by the eight differential analysis methods were transformed into ranks and plotted for pairwise comparison. Variants of the two components of Cepo—rCV and rzprop—were included for comparison. Spearman’s correlation was used to calculate the correlation between the ranks. ***P < 0.001. The scatter points are coloured by the proportion of zeros found in the gene expression across all cells.

**Supplementary Figure 5.**
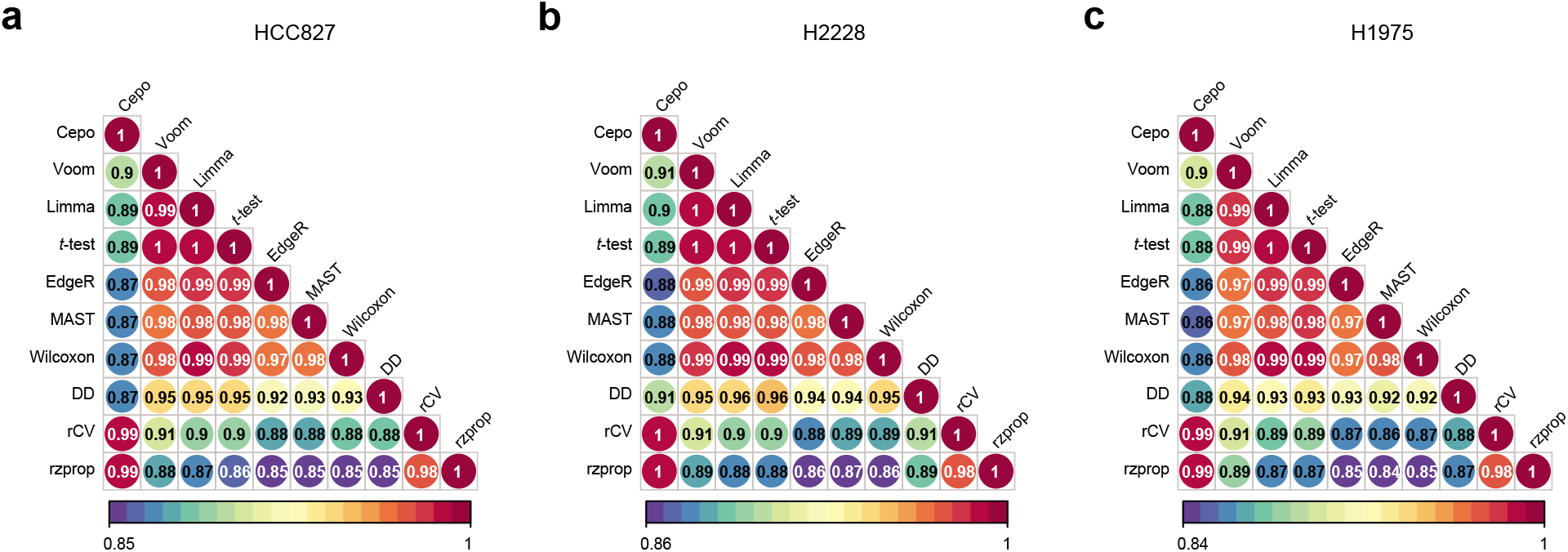
Correlation plot of ranked differential statistics for lung adenocarcinoma cell lines. Correlation plot showing the pairwise Spearman’s correlation between the differential statistics derived from the eight benchmarked differential analysis methods and the two components of Cepo, rCV and rzprop, for **(a)** HCC827, **(b)** H2228 and **(c)** H1975.

**Supplementary Figure 6.**
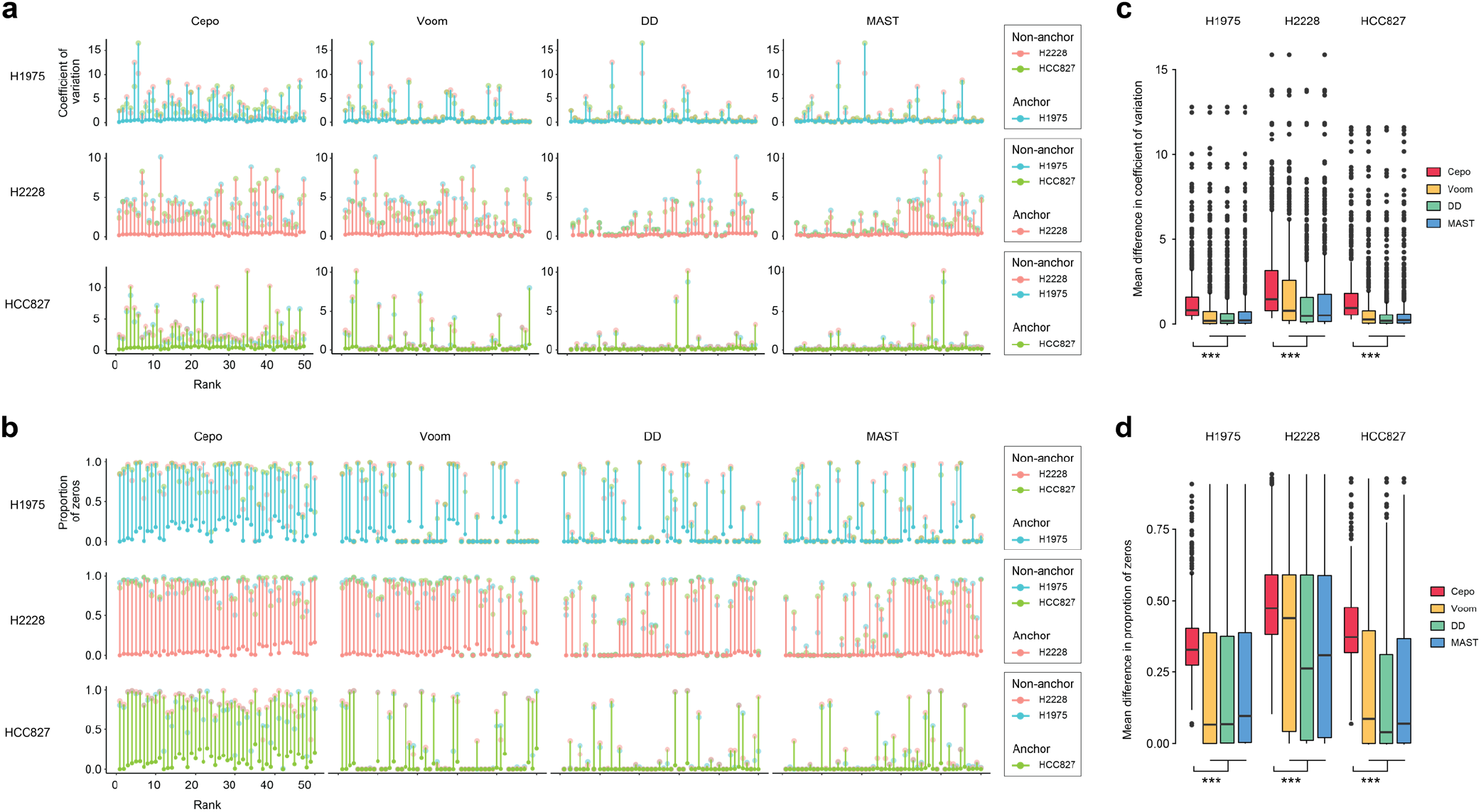
Differential patterns of the two components of Cepo—coefficient of variation and proportion of zeros—between cell types. Scatter plot of **(a)** the coefficient of variation (y-axis) and **(b)** proportion of zeros (y-axis) of top 50 differential analysis genes identified for each cell type (rows) by each method (columns). The genes are ordered by rank (x-axis), where a lower rank denotes a higher score. In each plot, the coefficient of variation and proportion of zeros of the genes were calculated and plotted by cell type, and vertical lines were drawn connecting the anchor point to the furthest nonanchor point, coloured using the anchor cell-type colour label. Boxplot of mean difference in **(c)** coefficient of variation and **(d)** proportion of zeros between the anchor cell type and the non-anchor cell types. **P < 0.01; ***P < 0.001, two-sided *t*-test.

**Supplementary Figure 7.**
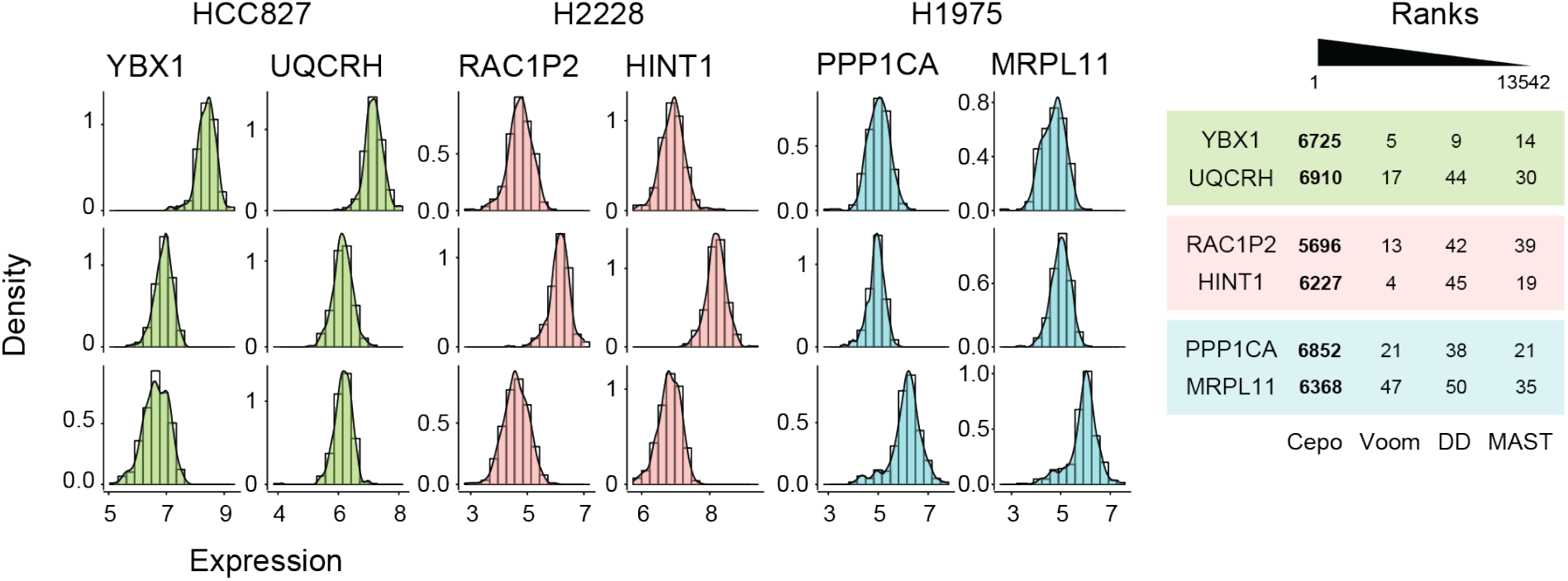
Distribution of top ranked DE genes with low differential stability. Distribution of example DE genes with low differential stability for each cell type. The ranking of each gene in their respective cell types of interest by each method are tabulated. A lower rank denotes higher prioritization of the gene.

**Supplementary Figure 8.**
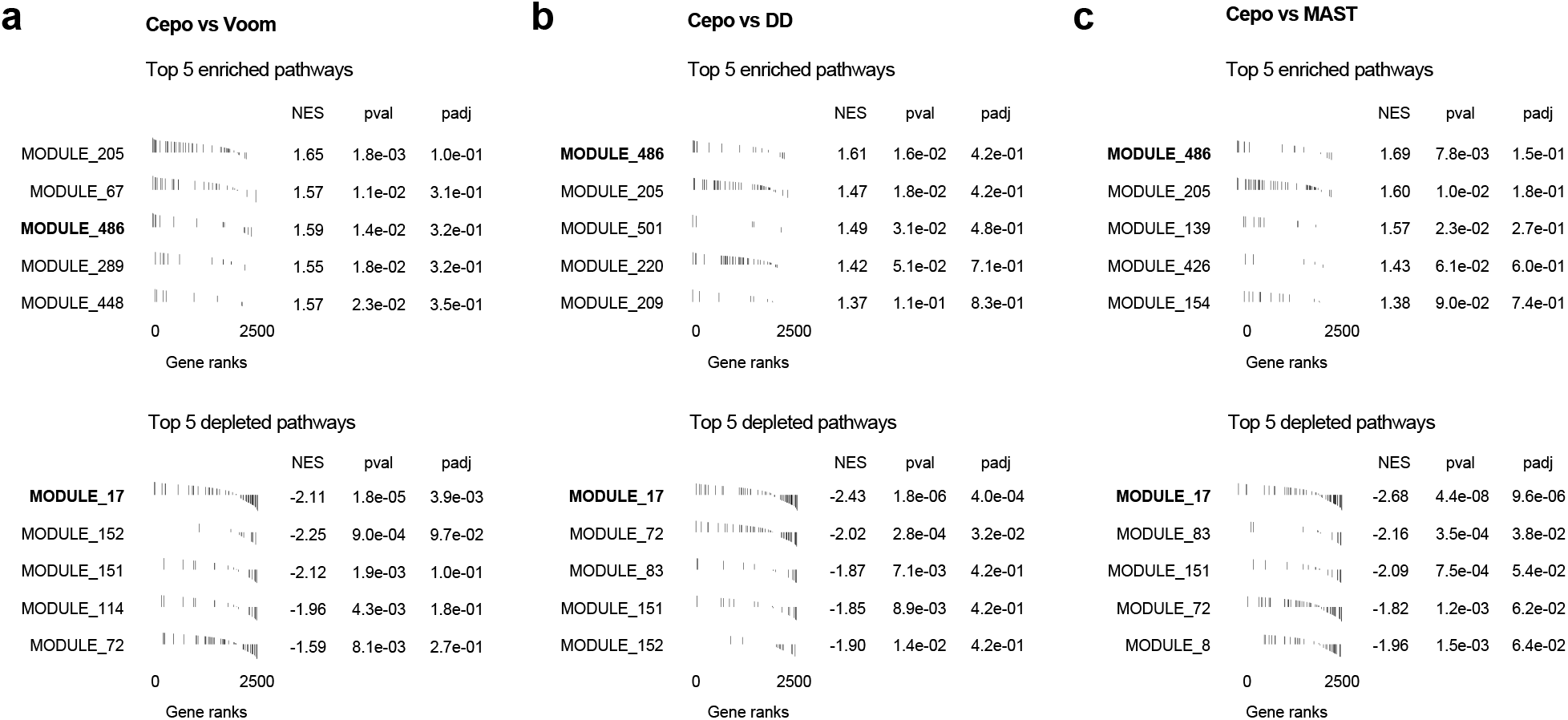
Visualisation of the top enriched and depleted pathways in the relative gene set enrichment analysis.

**Supplementary Figure 9.**
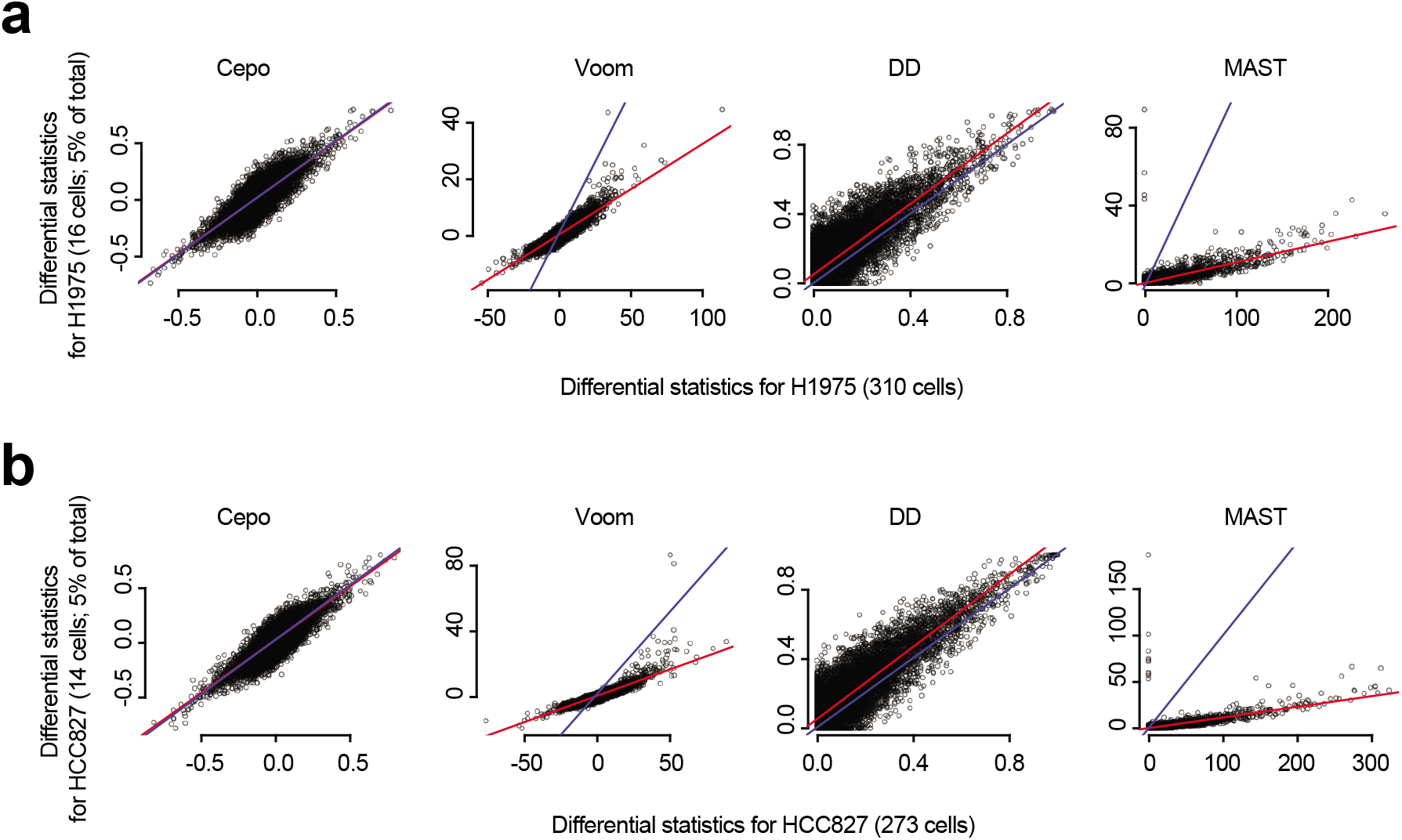
Scatter plot of differential analysis scores between rare and abundant cell types. Scatter plot of differential analysis scores from the full dataset (x-axis) and the rare cell type dataset (y-axis) for **(a)** H1975 and **(b)** HCC827. Rare cell types were artificially introduced by subsampling 5% of the total number of cells from each cell type. The red line denotes the best line of fit, and the blue line denotes *x* = *y*.

**Supplementary Figure 10.**
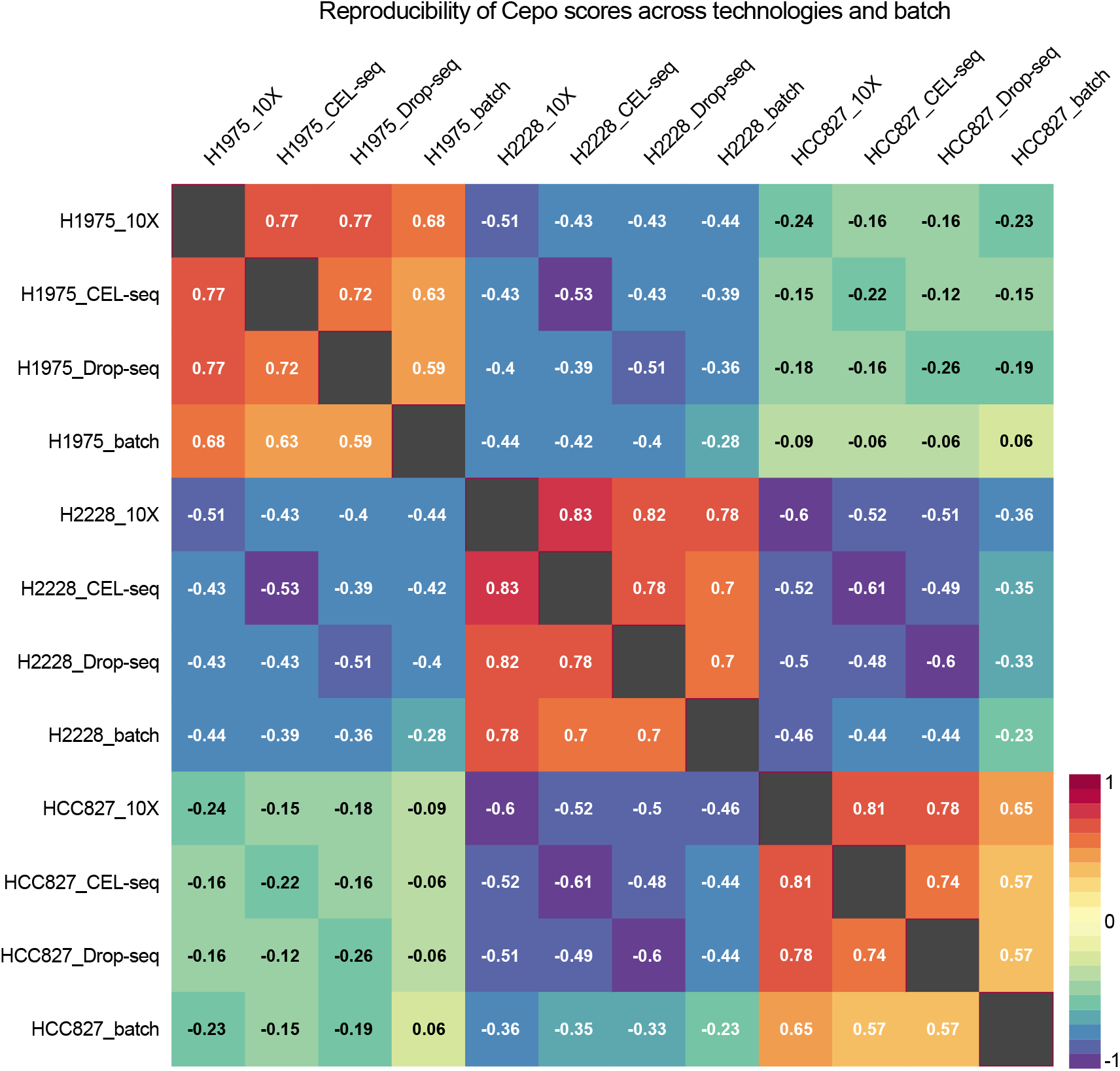
Reproducibility of Cepo across technology and batch. Correlation heatmap demonstrating the reproducibility of Cepo scores between three scRNA-seq technologies (10X Chromium, CEL-seq, and Drop-seq) and batch (10X Chromium). Spearman’s correlation was calculated on the differential analysis scores.

**Supplementary Figure 11.**
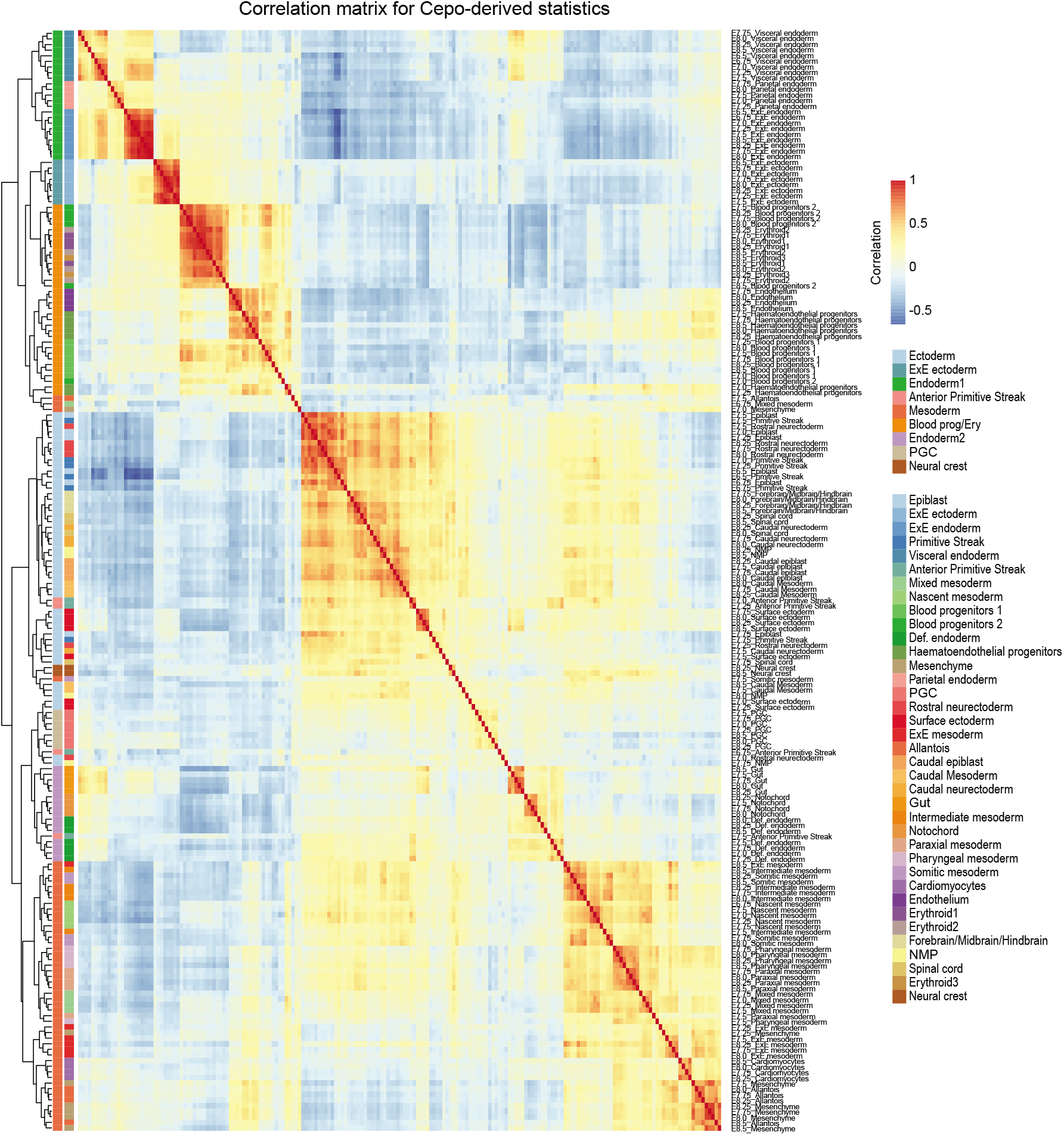
Correlation matrix of Cepo-derived statistics. Correlation matrix of the Cepo-derived statistics for the 196 samples (cell types resolved at different embryonic time points). Hierarchical clustering was performed on the correlation matrix. Row annotations denote the major lineages/cell-types and the original cell-type labels.

**Supplementary Figure 12.**
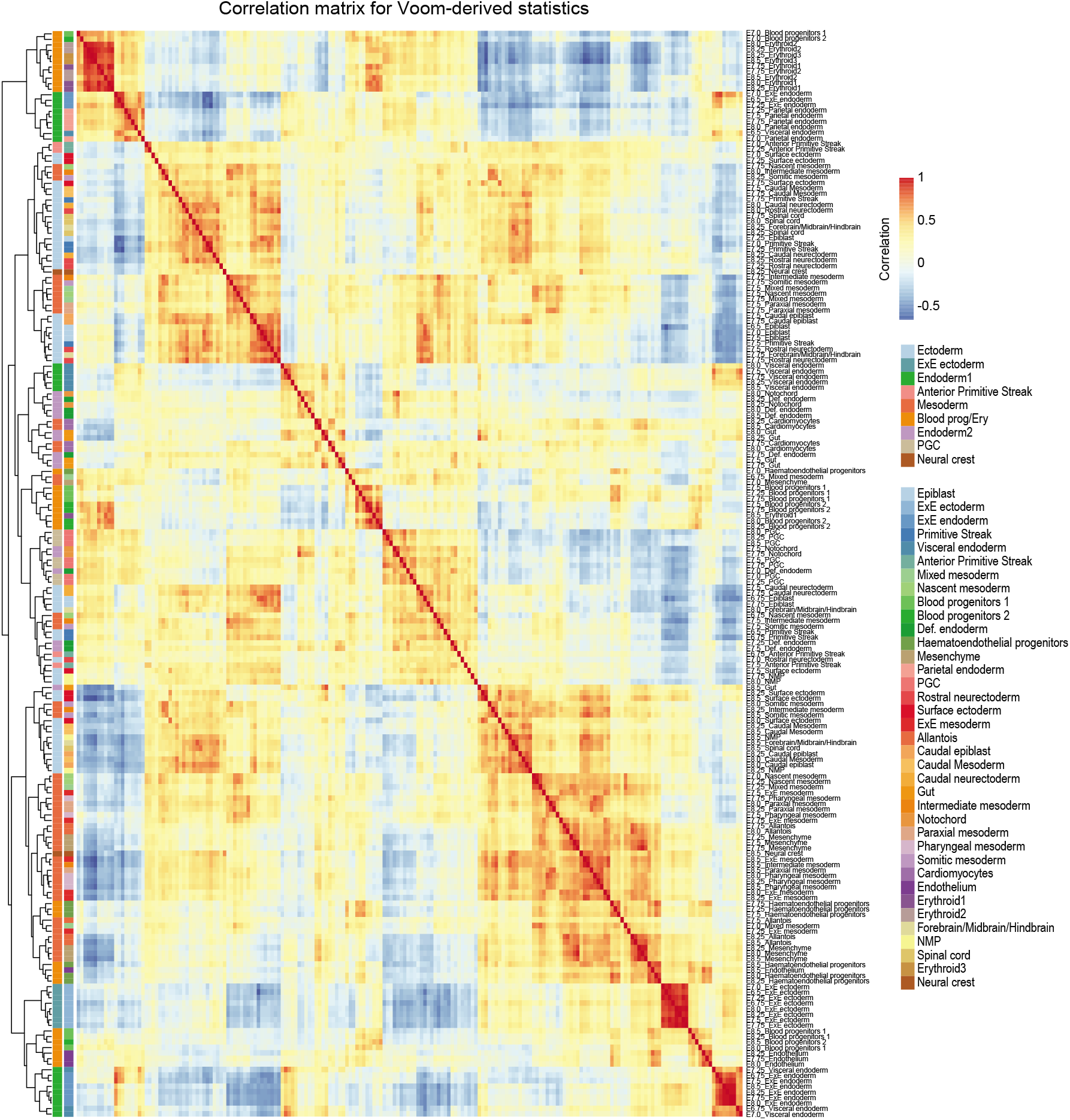
Correlation matrix of Voom-derived statistics. Correlation matrix of the Voom-derived statistics for the 196 samples (cell types resolved at different embryonic time points). Hierarchical clustering was performed on the correlation matrix. Row annotations denote the major lineages/cell-types and the original cell-type labels.

**Supplementary Figure 13.**
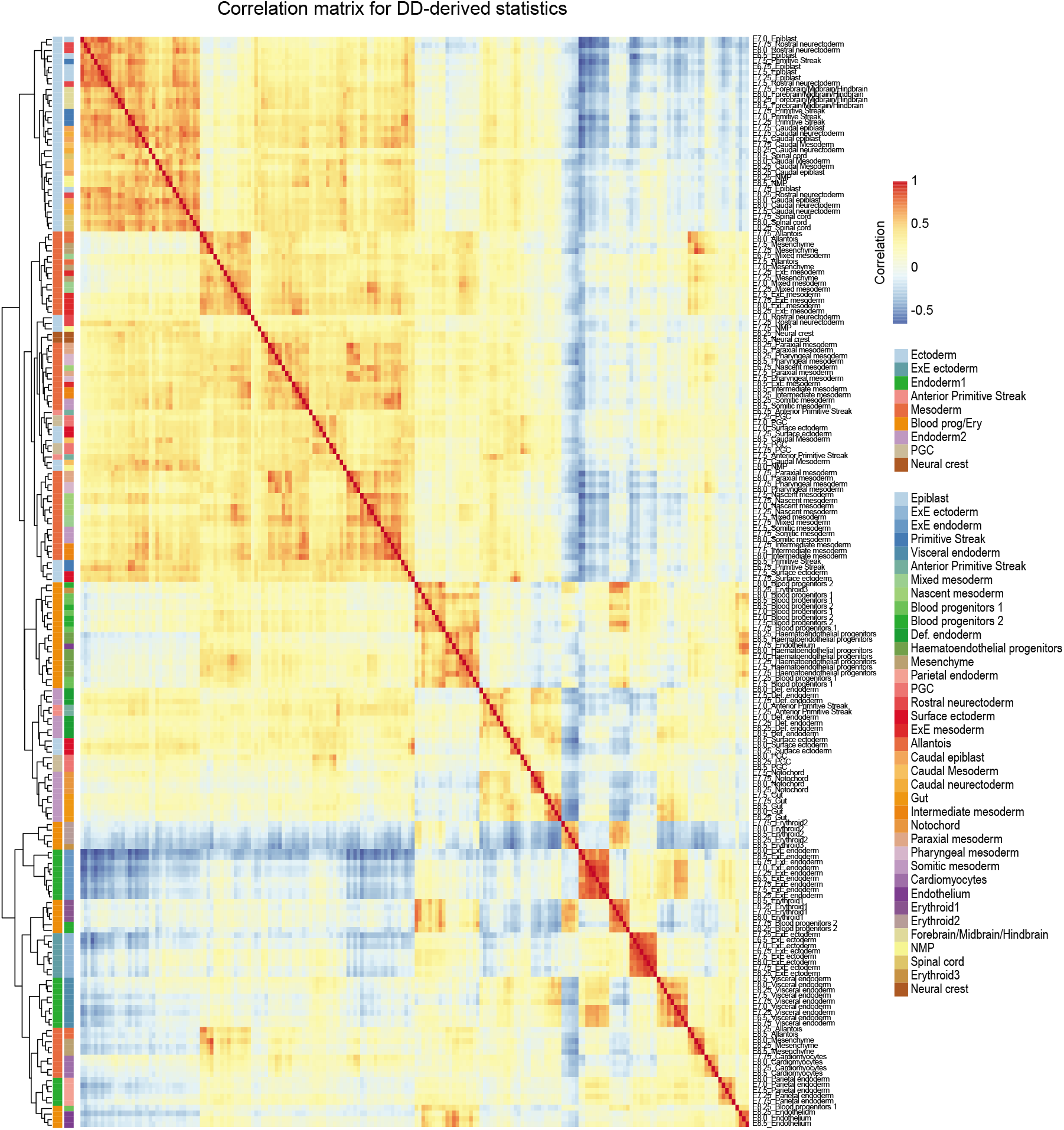
Correlation matrix of DD-derived statistics. Correlation matrix of the DD-derived statistics for the 196 samples (cell types resolved at different embryonic time points). Hierarchical clustering was performed on the correlation matrix. Row annotations denote the major lineages/cell-types and the original cell-type labels.

**Supplementary Figure 14.**
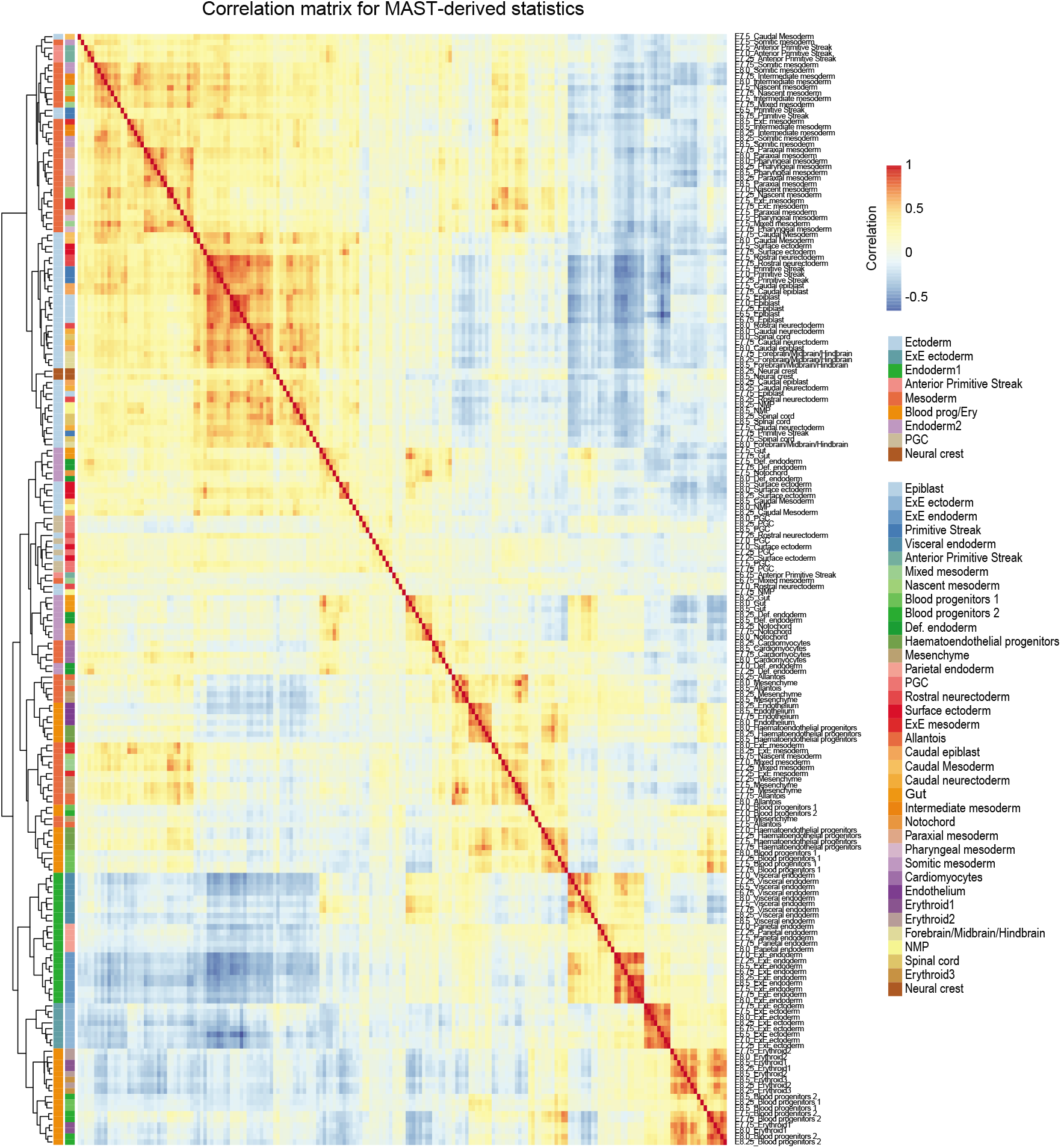
Correlation matrix of MAST-derived statistics. Correlation matrix of the MAST-derived statistics for the 196 samples (cell types resolved at different embryonic time points). Hierarchical clustering was performed on the correlation matrix. Row annotations denote the major lineages/cell-types and the original cell-type labels.

**Supplementary Figure 15.**
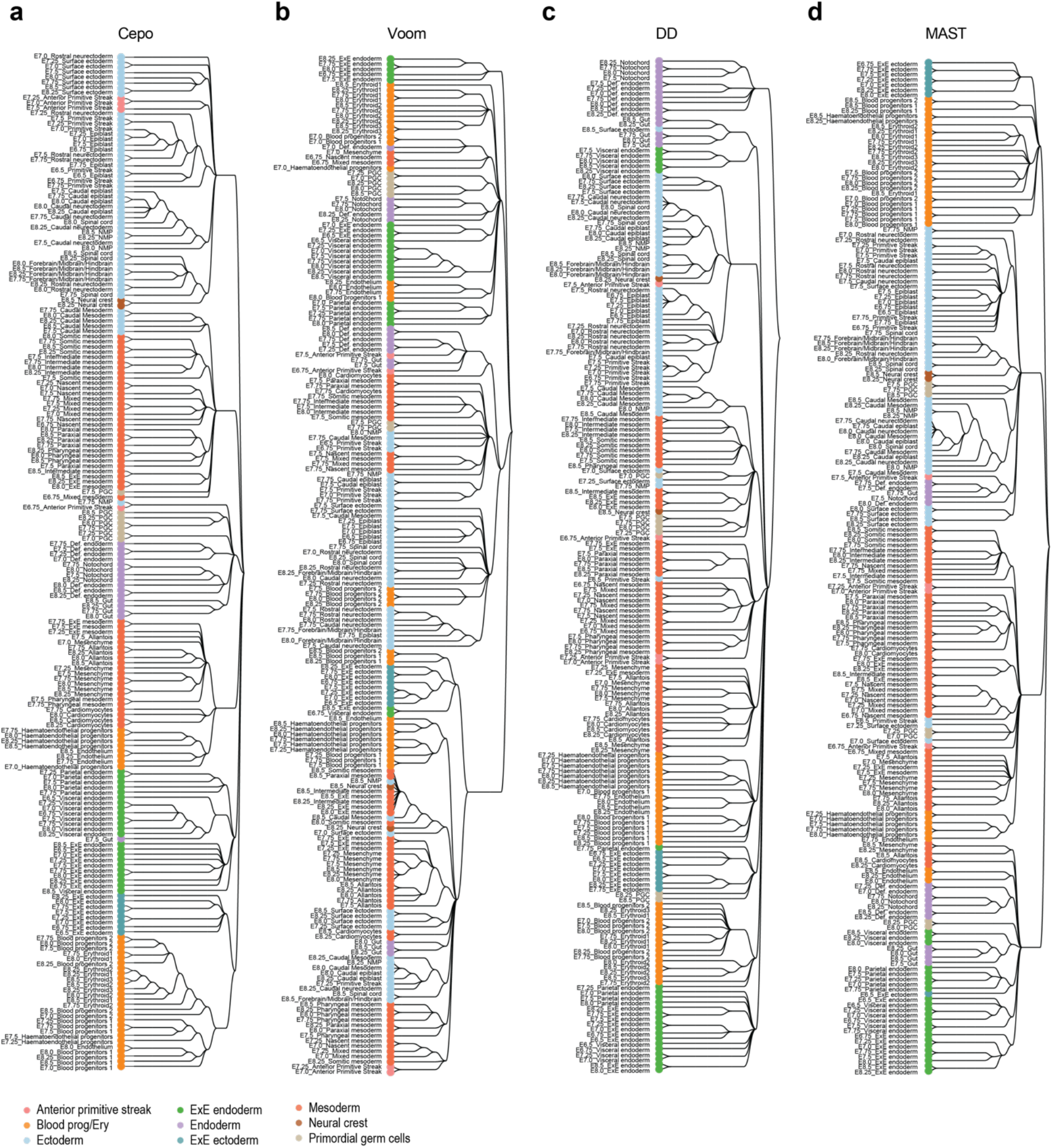
HOPACH clustering on differential statistics. HOPACH trees were generated by performing HOPACH clustering (kmax = 10) on the distance matrix of the 196 samples derived from the differential statistics of **(a)** Cepo, **(b)** Voom, **(c)** DD, and **(d)** MAST. The terminal nodes are coloured by the 9 major lineages/cell types.

**Supplementary Figure 16.**
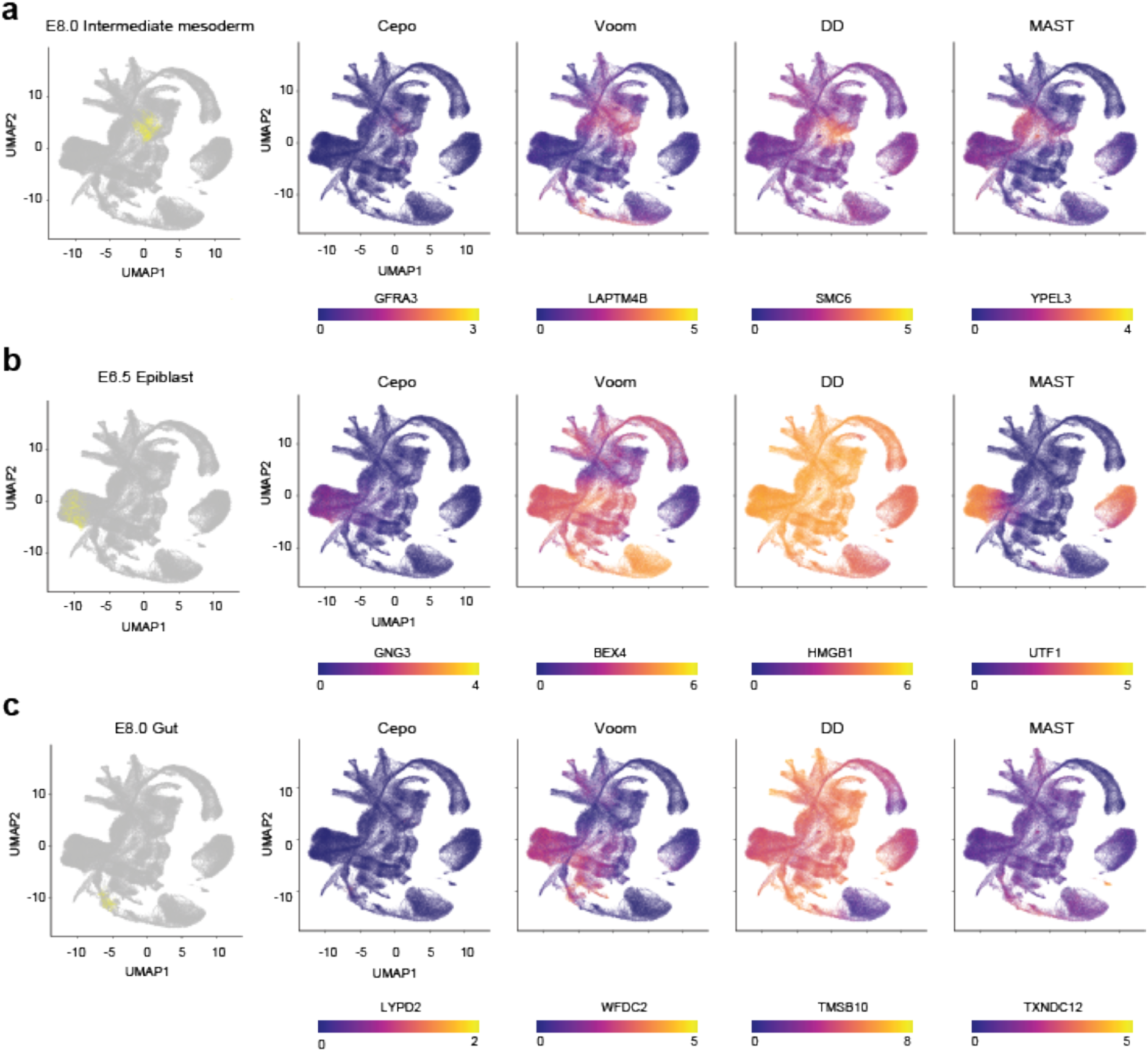
Gene expression of prioritised differential genes. Genes prioritised by each method were selected and its gene expression in the mouse embryogenesis atlas visualised on UMAP. As examples, genes identified for **(a)** E8.0 intermediate mesoderm, **(b)** E6.5 epiblast, and **(c)** E8.0 gut were highlighted for each method (columns).

**Supplementary Figure 17.**
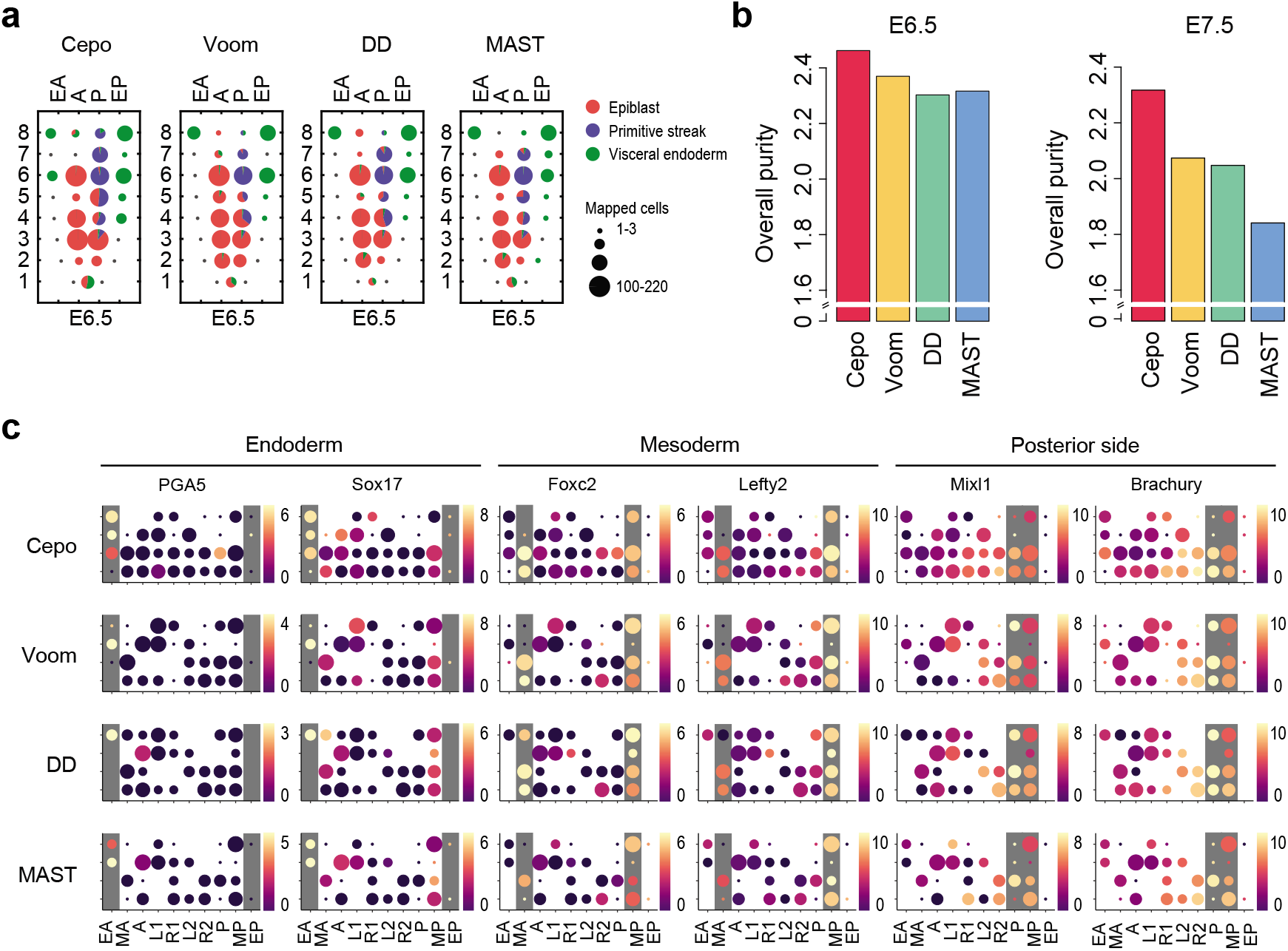
Spatial mapping of single cells onto the embryo. **(a)** Spatial mapping of E6.5 single cells from Smart-seq2 scRNA-seq dataset onto the E6.5 embryo (n = 31 spatial locations). Each dot denotes a pie chart that shows the proportion of cell types mapped to the location. The cell-type labels were defined as in the original publication. The size of each pie chart represents the number of cells mapped to each location. **(b)** Bar plots illustrating the overall purity of cell types mapped to the spatial locations of E6.5 and E7.5 embryos. (**c)** Non-zero expression of spatial marker genes of the endoderm, mesoderm, and posterior embryo. For each spatial gene (columns), the expected geographical location of expression has been marked with a grey background.

**Supplementary Figure 18.**
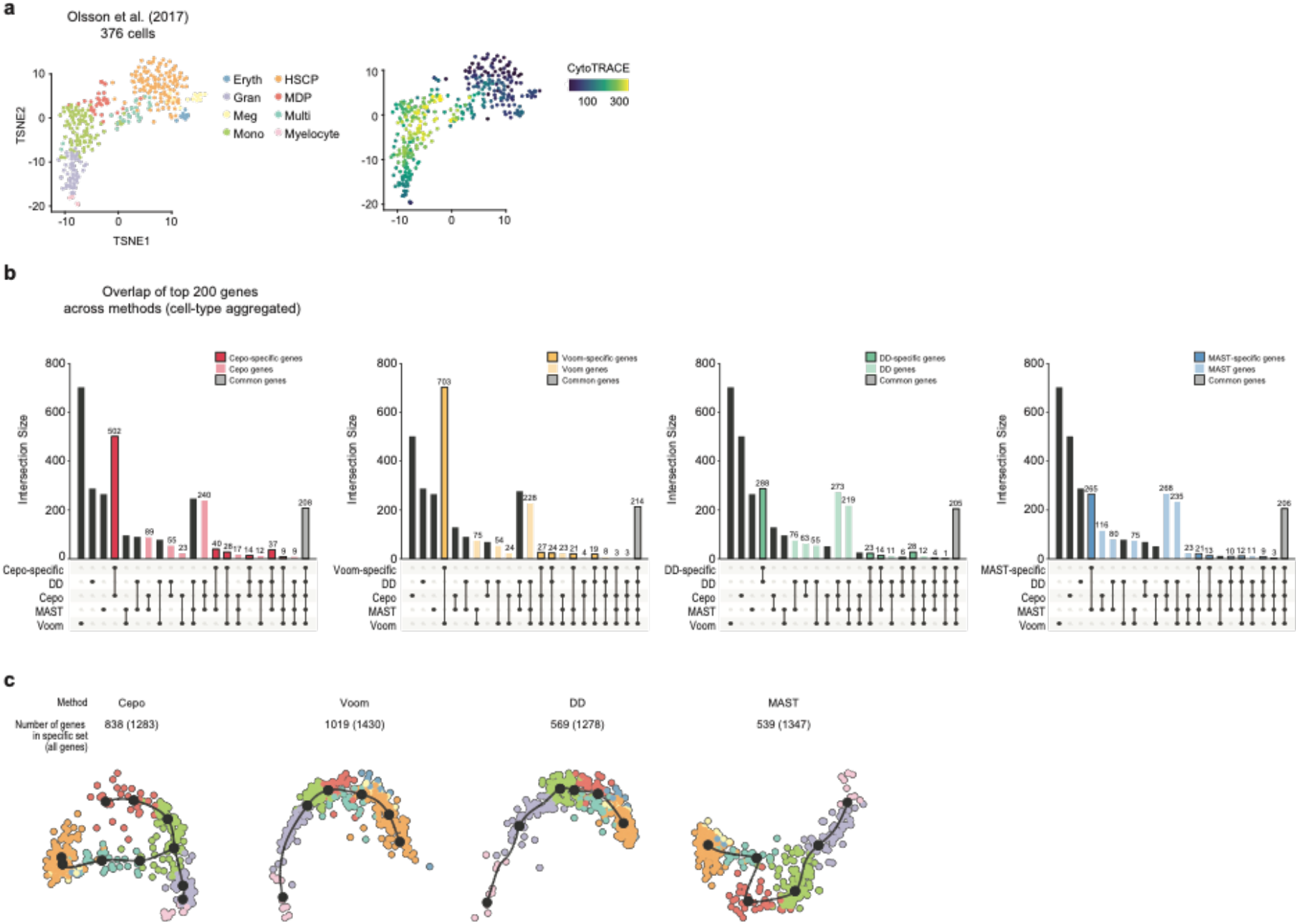
Trajectory inference on gene sets specifically identified by each method. **(a)** TSNE visualisation of the HSC differentiation scRNA-seq dataset. Individual points, denoting cells, are coloured by their original cell-type annotations (left) or by CytoTRACE-calculated pseudotime (right). **(b)** Upset plot illustrating the overlap between the differential genes identified by each method and the gene set specific to each method. The specific-gene set consists of genes that have been specifically identified by the method of interest to be among the top 200 genes for each cell type (dark-coloured bars) and genes that have been commonly identified by all methods (greycoloured bars). **(c)** Trajectory inference of hematopoietic stem cell differentiation scRNA-seq dataset using genes specifically identified by each differential analysis method (including the core common genes). For each method, the union of 200 genes per cell type were used prior to filtering of nonspecific genes. The multi-spanning tree method was used to generate the trajectory.

**Supplementary Figure 19.**
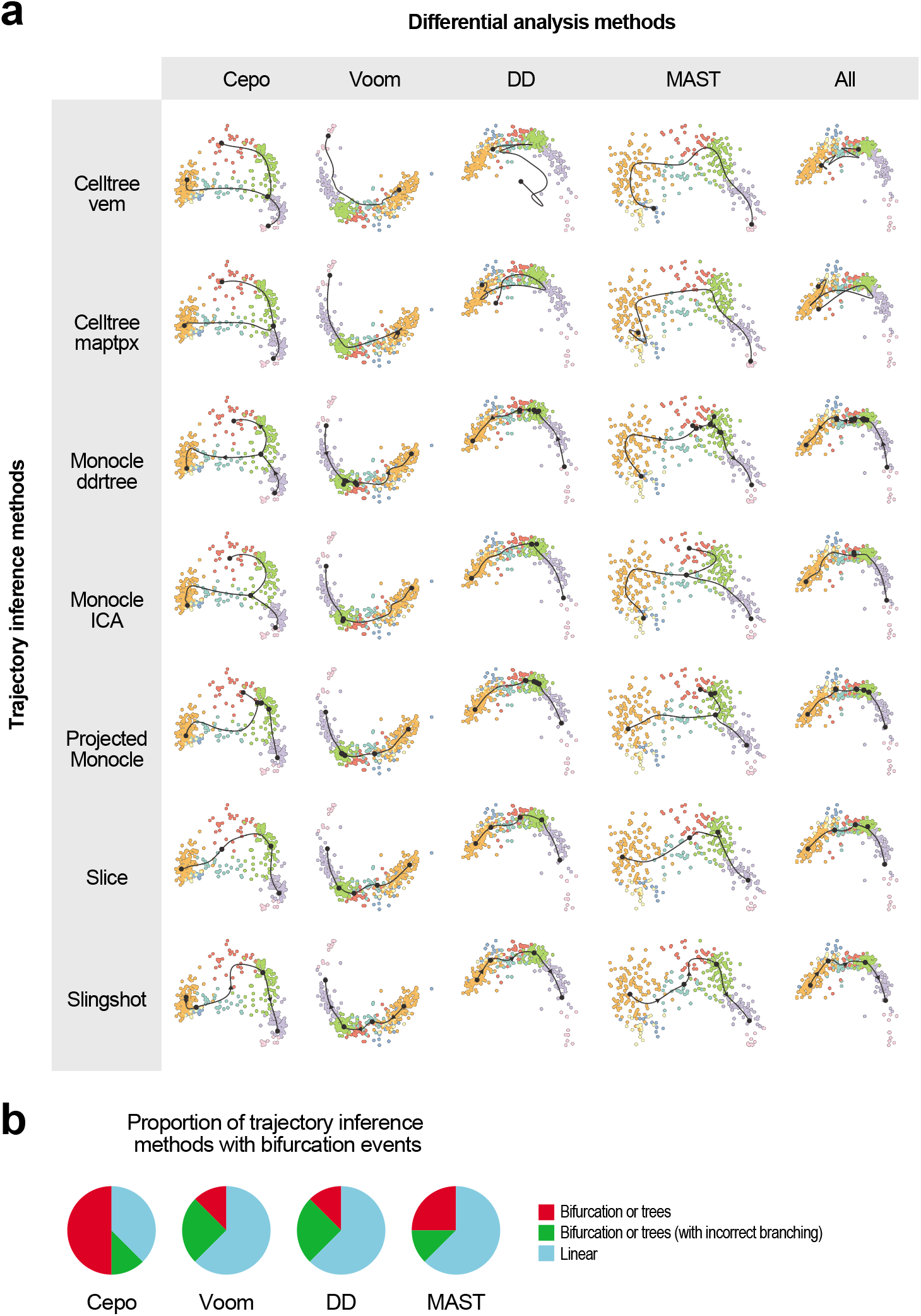
Trajectory inference using multiple inference tools. **(a)** Trajectories generated for the HSCP differentiation scRNA-seq dataset with 8 trajectory inference tools. For each cell type, the top 200 genes per cell type were selected, aggregated and then used in the trajectory analysis. To project the computed trajectory, the multi-spanning tree method for dimensionality reduction. Cells are coloured by cell type. **(b)** Pie charts showing the count of trajectory inference methods that are able detect a bifurcation event, where the terminal nodes encompass cell types of the two lineages, monocytes and granulocytes. A total of eight trajectory inference methods, as in **(a)** (Online Methods), and the top 200 genes per cell type were used to build the trajectories.

**Supplementary Figure 20.**
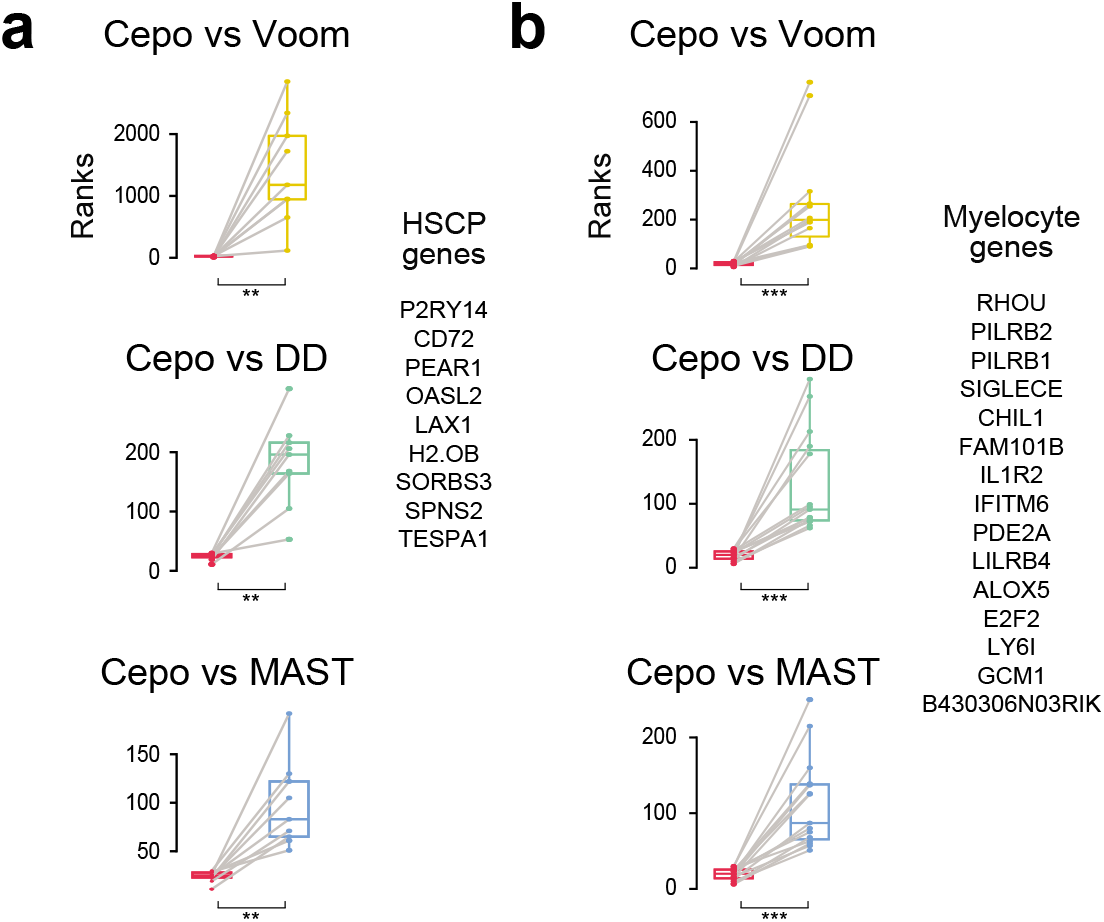
Paired boxplot of Cepo-prioritised genes. Paired boxplots of ranked differential statistics of select genes associated with **(a)** HSCPs and **(b)** myelocytes between Cepo and other differential analysis methods. **P < 0.01; ***P < 0.001, two-sided paired Wilcoxon test.

## Notes

### Competing Interest Statement

The authors have declared no competing interest.

## References

1. Kotliar, D. et al. Identifying gene expression programs of cell-type identity and cellular activity with single-cell RNA-Seq. Elife (2019). doi:10.7554/eLife.43803

2. Morris, S. A. The evolving concept of cell identity in the single cell era. Dev. (2019). doi:10.1242/dev.169748

3. Wagner, A., Regev, A. & Yosef, N. Revealing the vectors of cellular identity with single-cell genomics. Nature Biotechnology (2016). doi:10.1038/nbt.3711

4. Wang, T., Li, B., Nelson, C. E. & Nabavi, S. Comparative analysis of differential gene expression analysis tools for single-cell RNA sequencing data. BMC Bioinformatics (2019). doi:10.1186/s12859-019-2599-6

5. Soneson, C. & Robinson, M. D. Bias, robustness and scalability in single-cell differential expression analysis. Nat. Methods (2018). doi:10.1038/nmeth.4612

6. Korthauer, K. D. et al. A statistical approach for identifying differential distributions in single-cell RNA-seq experiments. Genome Biol. (2016). doi:10.1186/s13059-016-1077-y

7. Law, C. W., Chen, Y., Shi, W. & Smyth, G. K. Voom: Precision weights unlock linear model analysis tools for RNA-seq read counts. Genome Biol. (2014). doi:10.1186/gb-2014-15-2-r29

8. Robinson, M. D., McCarthy, D. J. & Smyth, G. K. edgeR: A Bioconductor package for differential expression analysis of digital gene expression data. Bioinformatics (2009). doi:10.1093/bioinformatics/btp616

9. Finak, G. et al. MAST: A flexible statistical framework for assessing transcriptional changes and characterizing heterogeneity in single-cell RNA sequencing data. Genome Biol. (2015). doi:10.1186/s13059-015-0844-5

10. Tian, L. et al. Benchmarking single cell RNA-sequencing analysis pipelines using mixture control experiments. Nat. Methods (2019). doi:10.1038/s41592-019-0425-8

11. Segal, E., Friedman, N., Koller, D. & Regev, A. A module map showing conditional activity of expression modules in cancer. Nat. Genet. (2004). doi:10.1038/ng1434

12. Cao, J. et al. A human cell atlas of fetal gene expression. Science (2020). doi:10.1126/science.aba7721

13. Pijuan-Sala, B. et al. A single-cell molecular map of mouse gastrulation and early organogenesis. Nature (2019). doi:10.1038/s41586-019-0933-9

14. Argelaguet, R. et al. Multi-omics profiling of mouse gastrulation at single-cell resolution. Nature (2019). doi:10.1038/s41586-019-1825-8

15. Tyser, R. et al. A spatially resolved single cell atlas of human gastrulation. bioRxiv (2020). doi:10.1101/2020.07.21.213512

16. Peng, G. et al. Molecular architecture of lineage allocation and tissue organization in early mouse embryo. Nature (2019). doi:10.1038/s41586-019-1469-8

17. Akashi, K., Traver, D., Miyamoto, T. & Weissman, I. L. A clonogenic common myeloid progenitor that gives rise to all myeloid lineages. Nature 404, 193–197 (2000).

18. Weinreb, C., Rodriguez-Fraticelli, A., Camargo, F. D. & Klein, A. M. Lineage tracing on transcriptional landscapes links state to fate during differentiation. Science (80-.). (2020). doi:10.1126/science.aaw3381

19. Olsson, A. et al. Single-cell analysis of mixed-lineage states leading to a binary cell fate choice. Nature (2016). doi:10.1038/nature19348

20. Ryzhov, S. et al. ERBB signaling attenuates proinflammatory activation of nonclassical monocytes. Am. J. Physiol. - Hear. Circ. Physiol. (2017). doi:10.1152/ajpheart.00486.2016

## References

1. Tian, L. et al. Benchmarking single cell RNA-sequencing analysis pipelines using mixture control experiments. Nat. Methods (2019). doi:10.1038/s41592-019-0425-8

2. Lun, A. T. L., Bach, K. & Marioni, J. C. Pooling across cells to normalize single-cell RNA sequencing data with many zero counts. Genome Biol. 17, 75 (2016).

3. Pijuan-Sala, B. et al. A single-cell molecular map of mouse gastrulation and early organogenesis. Nature (2019). doi:10.1038/s41586-019-0933-9

4. Argelaguet, R. et al. Multi-omics profiling of mouse gastrulation at single-cell resolution. Nature (2019). doi:10.1038/s41586-019-1825-8

5. Clark, S. J. et al. ScNMT-seq enables joint profiling of chromatin accessibility DNA methylation and transcription in single cells e. Nat. Commun. (2018). doi:10.1038/s41467-018-03149-4

6. Tyser, R. et al. A spatially resolved single cell atlas of human gastrulation. bioRxiv (2020). doi:10.1101/2020.07.21.213512

7. Picelli, S. et al. Smart-seq2 for sensitive full-length transcriptome profiling in single cells. Nat. Methods 10, 1096–1100 (2013).

8. Olsson, A. et al. Single-cell analysis of mixed-lineage states leading to a binary cell fate choice. Nature (2016). doi:10.1038/nature19348

9. Cao, J. et al. A human cell atlas of fetal gene expression. Science (2020). doi:10.1126/science.aba7721

10. Cao, J. et al. Comprehensive single-cell transcriptional profiling of a multicellular organism. Science (80-.). (2017). doi:10.1126/science.aam8940

11. Peng, G. et al. Molecular architecture of lineage allocation and tissue organization in early mouse embryo. Nature (2019). doi:10.1038/s41586-019-1469-8

12. Peng, G. et al. Spatial Transcriptome for the Molecular Annotation of Lineage Fates and Cell Identity in Mid-gastrula Mouse Embryo. Dev. Cell (2016). doi:10.1016/j.devcel.2016.02.020

13. McCarthy, D. J., Campbell, K. R., Lun, A. T. L. & Wills, Q. F. Scater: Pre-processing, quality control, normalization and visualization of single-cell RNA-seq data in R. Bioinformatics 33, 1179–1186 (2017).

14. Wang, T., Li, B., Nelson, C. E. & Nabavi, S. Comparative analysis of differential gene expression analysis tools for single-cell RNA sequencing data. BMC Bioinformatics (2019). doi:10.1186/s12859-019-2599-6

15. Soneson, C. & Robinson, M. D. Bias, robustness and scalability in single-cell differential expression analysis. Nat. Methods (2018). doi:10.1038/nmeth.4612

16. Korthauer, K. D. et al. A statistical approach for identifying differential distributions in single-cell RNA-seq experiments. Genome Biol. (2016). doi:10.1186/s13059-016-1077-y

17. Lin, Y. et al. Evaluating stably expressed genes in single cells. Gigascience 8, (2019).

18. Finak, G. et al. MAST: A flexible statistical framework for assessing transcriptional changes and characterizing heterogeneity in single-cell RNA sequencing data. Genome Biol. (2015). doi:10.1186/s13059-015-0844-5

19. Law, C. W., Chen, Y., Shi, W. & Smyth, G. K. voom: precision weights unlock linear model analysis tools for RNA-seq read counts. Genome Biol. 15, R29 (2014).

20. Robinson, M. D., McCarthy, D. J. & Smyth, G. K. edgeR: A Bioconductor package for differential expression analysis of digital gene expression data. Bioinformatics (2009). doi:10.1093/bioinformatics/btp616

21. Massey, F. J. The Kolmogorov-Smirnov Test for Goodness of Fit. J. Am. Stat. Assoc. (1951). doi:10.1080/01621459.1951.10500769

22. Benjamini, Y. & Hochberg, Y. Controlling the False Discovery Rate: A Practical and Powerful Approach to Multiple Testing. J. R. Stat. Soc. Ser. B (1995). doi:10.1111/j.2517-6161.1995.tb02031.x

23. Zappia, L., Phipson, B. & Oshlack, A. Splatter: Simulation of single-cell RNA sequencing data. Genome Biol. (2017). doi:10.1186/s13059-017-1305-0

24. Kuhn, M. & Vaughan, D. yardstick: Tidy Characterizations of Model Performance. (2020).

25. Pages, H. HDF5Array: HDF5 backend for DelayedArray objects. (2020).

26. Jindal, A., Gupta, P., Jayadeva & Sengupta, D. Discovery of rare cells from voluminous single cell expression data. Nat. Commun. (2018). doi:10.1038/s41467-018-07234-6

27. Su, S. et al. CellBench: R/Bioconductor software for comparing single-cell RNA-seq analysis methods. Bioinformatics (2020). doi:10.1093/bioinformatics/btz889

28. van der Laan, M. J. & Pollard, K. S. A new algorithm for hybrid hierarchical clustering with visualization and the bootstrap. J. Stat. Plan. Inference (2003). doi:10.1016/S0378-3758(02)00388-9

29. Kolde, R. pheatmap: Pretty Heatmaps. R package version 1.0.12. R package version 1.0.8 (2015).

30. Lin, Y. et al. scClassify: sample size estimation and multiscale classification of cells using single and multiple reference. 1–16 (2020). doi:10.15252/msb.20199389

31. Saelens, W., Cannoodt, R., Todorov, H. & Saeys, Y. A comparison of single-cell trajectory inference methods. Nat. Biotechnol. (2019). doi:10.1038/s41587-019-0071-9

32. Street, K. et al. Slingshot: Cell lineage and pseudotime inference for single-cell transcriptomics. BMC Genomics (2018). doi:10.1186/s12864-018-4772-0

33. Trapnell, C. et al. The dynamics and regulators of cell fate decisions are revealed by pseudotemporal ordering of single cells. Nat. Biotechnol. 32, 381–6 (2014).

34. duVerle, D. A., Yotsukura, S., Nomura, S., Aburatani, H. & Tsuda, K. CellTree: An R/bioconductor package to infer the hierarchical structure of cell populations from single-cell RNA-seq data. BMC Bioinformatics (2016). doi:10.1186/s12859-016-1175-6

35. Taddy, M. A. On estimation and selection for topic models. in Journal of Machine Learning Research (2012).

36. Sergushichev, A. A. An algorithm for fast preranked gene set enrichment analysis using cumulative statistic calculation. bioRxiv (2016).

37. Liberzon, A. et al. Molecular signatures database (MSigDB) 3.0. Bioinformatics (2011). doi:10.1093/bioinformatics/btr260

38. Segal, E., Friedman, N., Koller, D. & Regev, A. A module map showing conditional activity of expression modules in cancer. Nat. Genet. (2004). doi:10.1038/ng1434

39. Yu, G., Wang, L. G., Han, Y. & He, Q. Y. ClusterProfiler: An R package for comparing biological themes among gene clusters. Omi. A J. Integr. Biol. (2012). doi:10.1089/omi.2011.0118

